# Deconstructing the contributions of heterogeneity to combination treatment of hormone sensitive breast cancer

**DOI:** 10.1101/2023.05.19.541369

**Authors:** Samantha Linn, Jenna A. Moore-Ott, Robyn Shuttleworth, Wenjing Zhang, Morgan Craig, Adrianne L. Jenner

## Abstract

Combination therapies are fundamental to cancer treatments, including in breast cancer the most common invasive malignancy in women. Breast cancer treatment is determined based on molecular subtypes, and since 2016, combination palbociclib and fulvestrant has been used to treat hormone receptor-positive breast cancer. However, the impact of heterogeneity of the tumour landscape and tumour composition dynamics on scheduling decisions remains poorly understood. To elucidate the contributions of variability at multiple scales to treatment outcomes in hormone receptor-positive breast cancer, we developed a simple mathematical model of two unique estrogen receptor positive (ER+) breast cancer cell types and their response to combination treatment with palbociclib and fulvestrant. We used this model to understand how the initial fraction of either cell type may impact the fraction remaining after treatment and examined how heterogeneity in pharmacokinetics and pharmacodynamics result in a large distribution of outcomes. Our results suggest that the pharmacokinetics and pharmacodynamics of fulvestrant were the major drivers of final tumour size and composition. We then leveraged our model to guide therapeutic scheduling of combination palbociclib and fulvestrant, demonstrating the use of mathematical modelling to improve our understanding of cancer biology and treatments.

## 1 Introduction

Breast cancer is the most common invasive malignancy in women with a one in eight chance a woman will develop breast cancer in their lifetime [21, 38]. Treatment of breast cancer requires a multifaceted approach combining surgery, radiation, neoadjuvant, and adjuvant treatments [7]. There are five molecular subtypes of breast cancer, each with a different combination of cancer cells that over- or under-express receptors of estrogen (ER+/-), progesterone (PR+/-), and HER2 (HER2+/-) [25]. Effective treatment of these varying subtypes of breast cancer requires a deep understanding of heterogeneity in their responses to the different treatment types; unfortunately, there is still no completely curative treatment for any subtype.

Combination therapy (i.e., combining two or more therapeutic agents) is a cornerstone of cancer therapy [19]. The major goal of combination therapy in oncology is to enhance the therapeutic efficacy of a single anti-cancer drug through co-administration with a synergistic or additive drug that targets key pathways [19]. For example, metformin, an agent used to treat type 2 diabetes, was found to increase the susceptibility of a p53 breast cancer cell line to therapeutic molecule tumour necrosis factor-related apoptosis inducing ligand (TRAIL) [32]. To that end, a combination therapy in breast cancer with recognised potential is palbociclib combined with fulvestrant [17, 34], approved in early 2016 by the FDA to treat hormone receptor-positive breast cancer [36].

Palbociclib (brand name Ibrance) is an orally available, highly selective inhibitor of cyclin-dependent kinase 4 and 6 (CDK4 and CDK6) [31, 33, 29]. CDK4/6 are critical mediators of the cellular transition into the S phase and are crucial for the initiation, growth, and survival of many cancer types [8]. As such, pharmacological inhibitors of CDK4/6 are rapidly becoming a new standard of care for patients with advanced hormone receptor-positive breast cancer. Palbociclib thus forces cells to stay in the G1 phase in lieu of undergoing cell division (Figure 1). Importantly, palbociclib does not induce apoptosis but instead halts cellular division. According to the US National Institutes of Health (NCT0300797914), patients have historically been on a 21-day-on, 7-day-off palbociclib schedule, though there were concerns that the off days were the reason behind worse patient outcomes [39]. A 5-day-on, 2-day-off schedule has thus far shown better health outcomes for the treatment of ER+ breast cancer [39, 13], though this study is still ongoing.

**Fig. 1.**
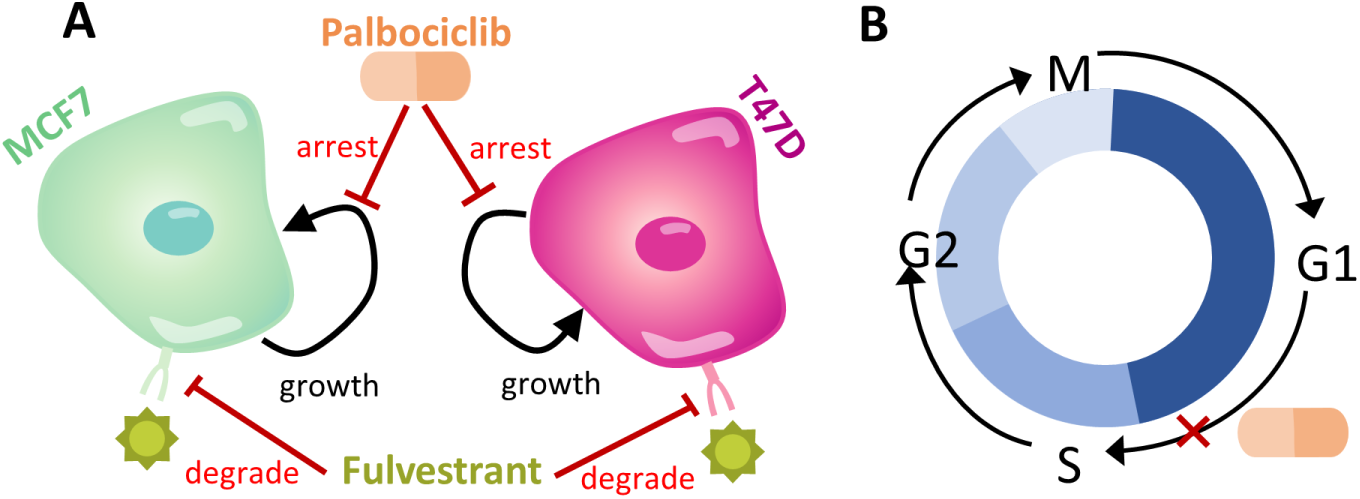
Schematic summarising the mathematical model of combination therapy on breast cancer cells. (A) Our model consists of pharmacokinetics and pharmacodynamics (PK/PD) of two drugs (palbociclib and fulvestrant), each with different mechanisms of action; palbociclib targets and arrests the cell in the cell cycle and fulvestrant degrades the estrogen receptor on cells, essentially causing cell death. To examine the impact of heterogeneity on tumour composition prior to, during, and at the end of treatment with this combination, we considered heterogeneous tumours composed of less aggressive and more aggressive cells. We parameterized these cells to in vitro data from two cell lines: MCF7 and T47D. (B) Schematic overview of the mechanism of action of palbociclib on cell cycle arrest.

Fulvestrant is a novel endocrine therapy for breast cancer which binds, blocks, and degrades the estrogen receptor, leading to complete inhibition of estrogen signaling through the ER [20, 11]. Through extensive pre-clinical and clinical trials, fulvestrant has demonstrated improved clinical efficacy compared to established endocrine agents [11]. Fulvestrant has been combined with several different classes of therapeutics, in particular CDK4/6 inhibitors [20]. The PALOMA-3 study investigated fulvestrant with palbociclib or placebo in both pre- and postmenopausal patients who had progressed on previous endocrine treatment. The trial demonstrated a substantial increase in progression free survival from 4.6 months to 9.5 months in the placebo compared to palbociclib arms [20]. The FLIPPER trial is a phase II study comparing fulvestrant and palbociclib with fulvestrant and placebo in the first-line metastatic setting, although this trial is ongoing.

While it can be challenging to fully capture the effects of heterogeneity on treatment outcomes experimentally and clinically, mathematical modelling is well-placed to provide insight into how cancer treatments are affected by multiple scales of heterogeneity. Previously, groups have used deterministic mathematical models to examine the combined treatment of breast cancer using palbociclib and AZD9496 [40]. For example, He et al. [9] used a mathematical model that captured the cell cycle and signalling pathways in response to endocrine therapy and CDK4/6 inhibition. Their model successfully predicted the combined effects of estrogen deprivation and palbociclib and was used to explore combination scheduling. Mathematical modelling studies can also be extended to a virtual or in silico clinical trial setting to account for variations in patient characteristics and comprehensively explore dosing regimens in ways that are clinically unfeasible [23, 24, 28, 1]. There are many examples of virtual clinical trials employed in cancer therapies [12, 43, 37, 3, 10] to this end, and this approach continues to gain traction within pharmaceutical and other applications [41, 2].

As heterogeneity can impact combination strategies aimed at CDK4/6 inhibition at multiple levels, we developed a simple mathematical model of two unique ER+ breast cancer cell types and their response to combination treatment with palbociclib and fulvestrant to understand how different sources of variation impact on this therapeutic approach. We examined in situ how co-culturing of heterogeneous cell types, specifically two commonly used breast cancer cell lines exhibiting different degrees of aggressivity, affects their response to treatment. Using our model, we next explored how interindividual variability in pharmacokinetics within a virtual breast cancer patient cohort affects treatment outcomes. Lastly, we used our integrated framework to establish how therapeutic scheduling determines treatment responses, providing insight into effective regimens using this combination treatment.

## 2 Methods

### 2.1 Mathematical model of breast cancer co-cultures and combination therapy

Below we detail the development of a mathematical model to capture the action of palbociclib and fulvestrant on a heterogeneous population of breast cancer cells. To capture and understand how intrinsic cell characteristics affect combination palbociclib and fulvestrant treatment, we considered co-cultures of MCF7 and T47D cell lines, two commonly used breast cancer cell lines that display different sensitivities to each drug, with T47D thought to be more aggressive (i.e., exhibit stronger/faster growth) than MCF7. A schematic overview of our model is provided in Figure 1.

#### 2.1.1 Palbociclib’s impact on cell growth

We first constructed a mathematical model describing the growth of a cell population under treatment by palbociclib, a drug whose effects were assumed to inhibit the cell cycle. To model these effects, we adopted a general inhibitory effects model given by

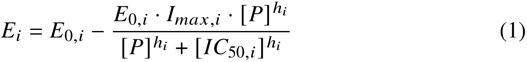

where *i* = *M*, *T* specifies the effect on cell line MCF7 and T47D respectively, *P* is the concentration of palbociclib at the tumour site, *E*_0,*i*_ denotes the baseline effect of the drug palbociclib on cell type *i*, *I*_*max*,*i*_ represents the maximal effect of the drug at high concentrations, ℎ_*i*_ is the Hill coefficient measuring the slope of the inhibitory curve for cell type *i*, and *IC*_50,*i*_ represents the drug concentration eliciting 50% of the maximal inhibition. This model formulation is regularly used to capture the effect of a drug on inhibiting a cell population [5].

As palbociclib arrests cells in the cell cycle, the general growth inhibition model for a population of cells of type *i*, i.e., *C*_*i*_ (*t*), inhibited by palbociclib is given by

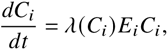

where *λ*(*C*_*i*_) is a function describing cell population growth in the absence of treatment (see Figure 1). As in vitro tumour growth is constrained by the availability of nutrients, space, etc., we modelled cell growth using the logistic growth law

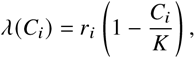

where *r*_*i*_ is the cell line specific proportionality constant and *K* is the total cell population carrying capacity in a given space. We chose logistic growth as this provided the most accurate fit to cell count measurements, however Gompertzian tumour growth also provided a close (but less accurate) fit to our data (Figure 10 and 11). Thus, the complete model of monoculture growth under treatment with palbociclib is given by

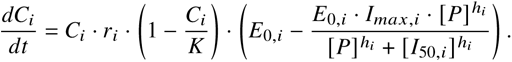

Typically, tumours are not homogeneous in nature and are comprised of a variety of different cell types. We accounted for this phenotypic heterogeneity by modelling both MCF7 and T47D cell types within a single tumour environment in co-culture. As mentioned above, while both MCF7 and T47D are ER+, they differ in their response to treatment. We therefore considered each to have separate parameters, with co-culture growth rates affected by the available space in the domain. In addition, since the effect parameters will depend on the drug being applied, we now update our pharmacodynamic variables to be drug specific. In other words, since parameters are cell line and drug specific, the specific combination is represented in their subscript as:

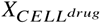

where *drug* is denoted by either *p* for palbociclib or *f* for fulvestrant and the cell line is denoted either by *M* for MCF7 or *T* for T47D.

For simplicity, and owing to the absence of data, we did not consider switching between tolerant and resistant types [4]. We assumed that the carrying capacity and growth of each cell type would be affected by the presence of the other cell type in the dish, thus modifying our logistic growth model. Therefore, to account for the impact of variable growth between each cell type, we included cell-specific carrying capacities in our model of cell growth (*λ*(*C*_*i*_) above). Our final model describing the change in population of two, indirectly interacting, cell types (MCF7 and T47D) is therefore given by

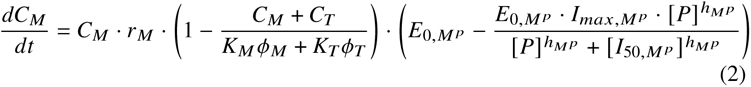

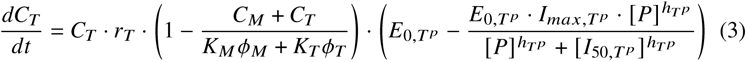

where *ϕ*_*i*_ is the volume fraction of cell type *i* for either type MCF7 and T47D, respectively, in the domain, and is calculated by *ϕ*_*i*_ = *C*_*i*_/(*C*_*M*_ +*C*_*T*_), with *ϕ*_*M*_ + *ϕ*_*T*_ = 1. The global carrying capacity in the domain is given by *K* = *K*_*M*_ *ϕ*_*M*_ + *K*_*T*_ *ϕ*_*T*_, where *K*_*i*_ is the individual carrying capacities for each cell type, MCF7 and T47D, respectively.

#### 2.1.2 Modelling the effects of fulvestrant on a heterogenous tumour

As fulvestrant degrades cells, we modelled its effect on the rates of decay for both the MCF7 and T47D cells. We updated the ordinary differential equation (ODE) system for the effect of palbociclib (Eqs. 2-3) to include a decay term for both cell populations, *d*_*i*_, that was then affected by the concentration of fulvestrant (*F*), using a modified version of the effect function in Eq. 1:

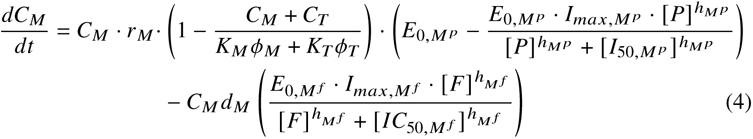

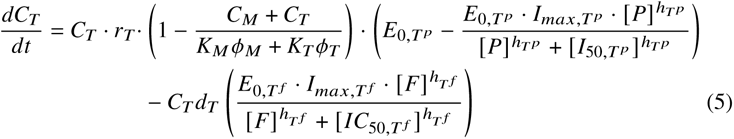

where *E*_0,*i f*_ was the basal effect of fulvestrant on cell type *i*, *I*_*max*,*i f*_ was the maximum effect of fulvestrant, ℎ_*i f*_ the Hill coefficient for fulvestrant and *IC*_50,*i f*_ was the half effect of fulvestrant, given *i* represented either *M* or *T* for cell type MCF7 or T47D respectively. Note, this modified effect function for fulvestrant is to capture that as the fulvestrant concentration *F* increases, the death rate increases. To determine the concentration of palbociclib and fulvestrant after administration, we introduced pharmacokinetic models parameterized from clinical pharmacokinetic (PK) studies for both drugs.

#### 2.1.3 Palbociclib and fulvestrant pharmacokinetic models

We used a linear two-compartment pharmacokinetic (PK) model with first-order absorption and absorption lag to model the dynamics of orally administered palbociclib,

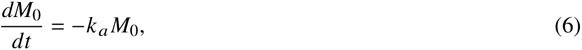

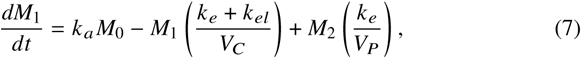

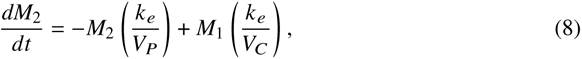

where *M*_0_, *M*_1_ and *M*_2_ are the palbociclib concentrations pre-absorption, in plasma, and in peripheral tissue, respectively. Further, *k*_*a*_ is the rate of absorption into plasma, *k*_*el*_ is the rate of linear elimination, *k*_*e*_ is the exchange rate between plasma and tissue, and *V*_*C*_ and *V*_*P*_ are the apparent plasma and peripheral tissue volumes, respectively. The concentration of palbociclib at the tumour site was then calculated by *P*(*t*) = *M*_1_(*t*)/*V*_*C*_ in Eqs. 4-5.

Based on data from 38 postmenopausal women with advanced breast cancer who received 250 mg doses of extended-release fulvestrant (in a single 5 mL intramuscular (IM) injection or two 2.5 mL IM injections [26]), a two-compartment PK model with zero-order administration and linear elimination was developed. The fulvestrant PK model is given by

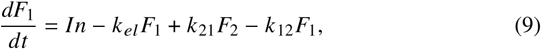

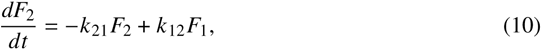

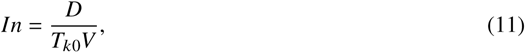

where *F*_1_ and *F*_2_ denote fulvestrant concentrations in the plasma and tissues, respectively, *In* represents the administered dose (here taken to be an intramuscular administration), *k*_*el*_ is the rate of linear elimination, *k*_12_ and *k*_21_ are transit rates between the plasma and tissue compartments, *D* represents the IM dose, *T*_*k*0_ the time for absorption, and *V* the volume of distribution. The concentration of fulvestrant at the tumour site was set as *F* (*t*) = *F*_1_(*t*) in Eq. 4-5.

### 2.2 Parameter estimation

#### 2.2.1 Estimating tumour growth parameters

Cell counting was performed in breast cancer cell lines MCF7 and T47D by Vijayaraghavan et al. [35]. Cells were plated in six-well plates and treated with indicated agents for 10 days. the medium was replaced every other day over the course of the experiment. Cells were then collected and counted using BioRad TC20 Automated Cell Counter on days 0, 3, 6 and 10 (see data in Figure 10 and 11). We estimated parameters governing cell growth by setting all drug concentrations to zero in our model Eqs. 4-5 and fitting the proliferation rate *r*_*i*_ and carrying capacity *K*_*i*_ to cell type i count data. Fitting was performed in Matlab [18] using the inbuilt non-linear least-squares fitting function *lsqnonlin*. The model was solved using ode45 and the trust-region-reflective algorithm with 1000 maximum function evaluations was chosen.

#### 2.2.2 Estimating drug effect parameters from cell viability assays

Cell viability measurements for MCF7 and T47D with palbociclib were measured by Vijayaraghavan et al. [35]. For these dose-response studies, cells were plated on a 96- well plate and treated with increasing concentrations (0.01-12 *μ*M) of palbociclib for 1, 2, 4, 6, or 8 days. The medium was replaced with drug-containing medium every other day. At completion of drug treatment, cultures were continued in drug-free medium until day 12 after which they were stained with 0.5% crystal violet solution. Values were normalized to those of their no treatment controls. We assumed that after 8 days of drug exposure, the drug effects were saturated. We fit the 8-day data (see Figure 12) and estimated the values of *E*_0,*ip*_, *I*_*max*,*ip*_, and *IC*_50,*ip*_ for each cell type *i* in Eqs. 4-5 by minimizing the least-squares error between the data and the inhibitory growth model using *lsqnonlin* in Matlab [18]. We additionally estimated the 95% confidence intervals for the parameters using the Jacobian returned from the *lsqnonlin* fit. All fitted parameters and their bounds are given in Figure 12 for both MCF7 and T47D cell lines.

In similar experiments, Nukatsuka et al. [22] measured MCF7 cell growth under varying fulvestrant concentrations. Measurements were calculated as means and standard deviation of cell growth relative to that of the control for three independent experiments. Lewis-Wambi et al. measured DNA (*μ*g/well) from T47D cells after treatment with fulvestrant [15]. Cells were seeded in 24-well dishes and after 24h were treated with varying drug concentration for 7 days. At the conclusion of the experiment, cells were harvested, and proliferation was assessed as cellular DNA mass (in micrograms per well). We assumed this as a proxy for cell viability relative to control. As with the palbociclib experiments from Vijayaraghavan et al. [35], we estimated the pharmacodynamic parameters in Eq. 4-5 by minimizing the least-squares error between the data and the inhibitory growth model using *lsqnonlin* in Matlab (see Figure 12).

#### 2.2.3 Estimating pharmacokinetic parameters

We used a nonlinear mixed-effects model in Monolix [16] to estimate parameters of the fulvestrant PK model in Eqs. 9 and 11. As the data reported in Robertson et al. [26] was pooled, we extracted the reported mean, and lower and upper bounds to estimate interindividual variability (IIV). We then fit the model in Eq. 9 - 11 to this data assuming lognormal distributions on parameters (Figure 13) subject to IIV according to

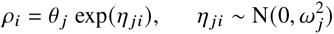

where *ρ*_*i*_ is the value of a given model parameter (e.g., *k*_*el*_, *k*_12_ etc.) for subject *i*, *θ*_*j*_ is the population mean, and *η* _*ji*_ represents the deviation from the mean (i.e., IIV) for the *i*-th individual. Estimated model parameters are presented in Tables 1 and 2.

**Table 1.** Estimated parameter values for the palbociclib population pharmacokinetic model.

| Parameter | Units | Fixed effects |  |
| --- | --- | --- | --- |
|  |  | Mean | Variance |
| $k_a$ | 1/hour | 0.617 | 0.484 |
| $k_{el}/F$ | L/hour | 48 | 0.0451 |
| $k_e/F$ | L/hour | 4.49 | 0.61 |
| $V_C/F$ | L | 1520 | 0.0185 |
| $V_P/F$ | L | 2780 | - |

**Table 2.**
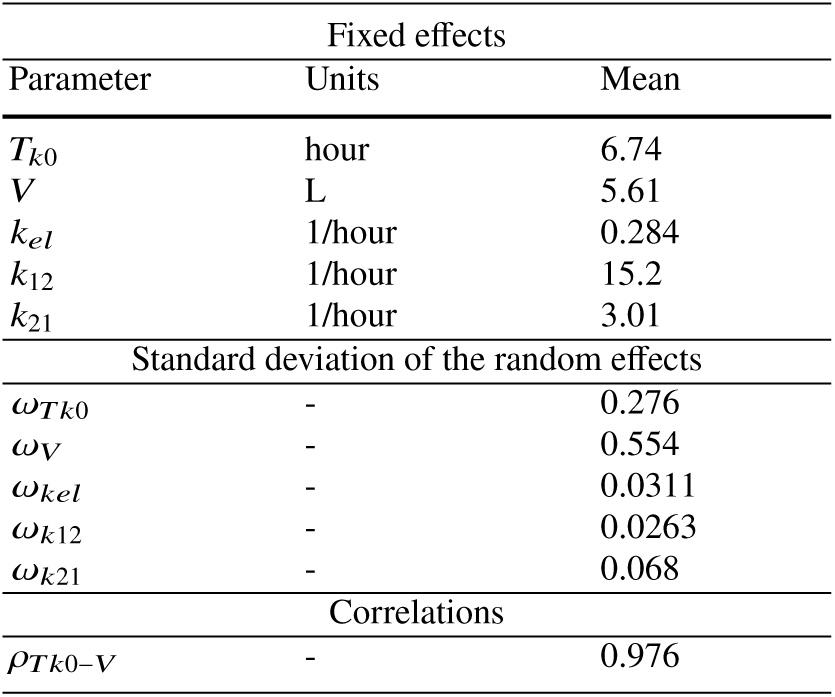
Estimated parameter values for the fulvestrant population pharmacokinetic model.

The model in Eqs. 6 - 8 is based on the clinical and theoretical work of Yu et al. [42] which described data from 26 advanced breast cancer patients who received palbociclib and letrozole on a three-weeks-on, one-week-off treatment regimen. The model parameters were taken from Yu et al. and used to simulate patient populations. We assumed lognormal distributions on parameters subject to interindividual variability using *ρ*_*i*_ above. Parameter values for fulvestrant were taken directly from Robertson et al. [26].

### 2.3 Generating heterogeneous pharmacokinetics and pharmacodynamics

#### 2.3.1 Pharmacokinetic parameters

We investigated palbociclib and fulvestrant individually to quantify each of their contributions to the effects of PK IIV on tumour growth. For palbociclib, we sampled *V*_*C*_, *V*_*P*_, *k*_*el*_, *k*_*e*_, and *k* _*a*_ from lognormal distributions according to the parameters in Table 1 to produce a virtual patient population. Similarly, for fulvestrant, we sampled *T*_*k*0_, *V*, *k*_*el*_, *k*_12_, and *k*_21_ from lognormal distributions according to the bestfit nonlinear mixed effects model determined by our parameter fitting (see Table 2) to generate virtual patients. In the case of each drug, by simulating the full model (with all other components’ parameters set to their average values), we selected only those virtual patients whose predicted trajectories were realistic (as confirmed by visual predictive check of their concentration time courses) before accepting them into our cohort. This left 500 virtual patients in the case of palbociclib and 438 for fulvestrant.

#### 2.3.2 Pharmacodynamic parameters

To investigate the effect of heterogeneity of the pharmacodynamics of palbociclib and fulvestrant, we generated 400 sets of parameter values by sampling *E*_0_, *I*_*max*_, ℎ and *IC*_50_ for each cell type-drug combination from the ranges established during parameter fitting (Figure 14). As each of these four parameters is drug and cell type specific, this gave 16 parameters to sample:

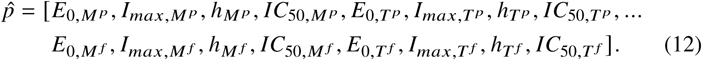

Parameters were sampled from a multivariate normal distribution with mean *μ* set to the fitted values in Table 3 for *p̂* and standard deviation *σ* determined from the confidence intervals (CI) returned for the fitted parameters and the formula

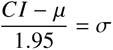

where 1.95 was chosen to return values in the 95% confidence interval. Any samples resulting in negative parameter values were discarded. The resulting distributions of parameters are provided in Figure 14.

**Table 3.** Table of fitting parameter values. Values in this table were obtained in fits in Figure 10 and Figure 11.

| Cell line | Variable | Description | Units | Fit | Bounds |  |
| --- | --- | --- | --- | --- | --- | --- |
|  |  |  |  |  | Upper | Lower |
| MCF7 | $r_M$ | Growth rate | 1/day | 0.6083 | 0.5390 | 0.6776 |
| | $k_M$ | Carrying capacity | cells | $7.61 \times 10^6$ | $3.98 \times 10^6$ | $11 \times 10^6$ |
| | $E_{0,M^P}$ | Palbo. initial no-drug effect | - | 98.9 | 96.1 | 101.7 |
| | $I_{max,M^P}$ | Palbo. max inhibition | - | 0.879 | 0.86 | 0.9 |
| | $h_{M^P}$ | Palbo. hill coefficient | - | 1.65 | 1.38 | 1.91 |
| | $IC_{50,M^P}$ | Palbo. half-effect | $\mu M$ | 0.67 | 0.61 | 0.73 |
| | $E_{0,M^f}$ | Fulv. initial no-drug effect | - | 96.3 | 95.4 | 97.2 |
| | $I_{max,M^f}$ | Fulv. max inhibition | - | 0.81 | 0.81 | 0.82 |
| | $h_{M^f}$ | Fulv. hill coefficient | - | 1.85 | 1.75 | 1.95 |
| | $IC_{50,M^f}$ | Fulv. half-effect | $\mu M$ | $4.4 \times 10^{-4}$ | $4.3 \times 10^{-4}$ | $4.5 \times 10^{-4}$ |
| T47D | $r_T$ | Growth rate | 1/day | 0.6726 | 0.6512 | 0.6941 |
| | $k_T$ | Carrying capacity | cells | $5.27 \times 10^6$ | $4.6 \times 10^6$ | $5.9 \times 10^6$ |
| | $E_{0,T^P}$ | Palbo. initial no-drug effect | - | 96.2 | 87.5 | 96.2 |
| | $I_{max,T^P}$ | Palbo. max inhibition | - | 0.95 | 0.79 | 1.11 |
| | $h_{T^P}$ | Palbo. hill coefficient | - | 1.02 | 0.53 | 1.52 |
| | $IC_{50,T^P}$ | Palbo. half-effect | $\mu M$ | 1.13 | 0.64 | 1.62 |
| | $E_{0,T^f}$ | Fulv. initial no-drug effect | - | 100 | 82.5 | 117.5 |
| | $I_{max,T^f}$ | Fulv. max inhibition | - | 0.84 | 0.74 | 0.95 |
| | $h_{T^f}$ | Fulv. hill coefficient | - | 1.85 | 0.2 | 1.32 |
| | $IC_{50,T^f}$ | Fulv. half-effect | $\mu M$ | $1.2 \times 10^{-4}$ | $-1 \times 10^{-4}$ | $2 \times 10^{-4}$ |

## 3 Results

### 3.1 Shorter treatment cycle reduces aggressive cell viabilities as compared to conventional schedule

We first set out to predict whether a shortened treatment cycle (i.e., 5 days on of palbociclib with 2 days of rest, repeated for 28 days) was a viable strategy as compared to a conventional 21 days on of palbociclib with 7 days of rest schedule. For this, we simulated the complete model in Eqs. 4-9 with mean values for both the palbociclib and fulvestrant pharmacokinetic models (Table 1 and 2) and pharmacodynamic effects model (Table 3). We considered only the case where the two cell types were present in equal fractions (i.e., *ϕ*_*i*_ = 0.5) with a total cell count of *C*_*M*,*i*_+*C*_*T*,*i*_ = 7×10^4^ cells. We called this an “average patient”. Left untreated, over the course of 28 days, unsurprisingly both cell lines were predicted to grow to the global carrying capacity *K* = *K*_*M*_ *ϕ*_*M*_ + *K*_*T*_ *ϕ*_*T*_ (see Figure 15).

We then introduced treatment to this average patient. We first simulated 125 mg of palbociclib daily for 21 days, with a period of rest for 7 days, with 125 mg of fulvestrant on days 1 and 15, consistent with current treatment schedules [42] (Figure 2). Our model predicted the resulting viability at the end of treatment to be 0.39 (MCF7) and 0.51 (T47D). Here, cell viability was determined by comparing, for the same parameters, treatment outcomes to the untreated scenario, i.e., the viability of each cell line was calculated by comparing the total cells at the end of treatment to the total number of cells under no treatment. Repeating this strategy for a treatment course of 125 mg of palbociclib for 5 days with 2 days of rest over a period of 28 days, with 125 mg of fulvestrant administered on days 1 and 15, we found viabilities after 28 days of 0.38 (MCF7) and 0.47 (T47D), respectively (Figure 2). Notably, this change in treatment schedule was predicted to somewhat lower the viability of T47D, which is the more aggressive cell type.

**Fig. 2.**
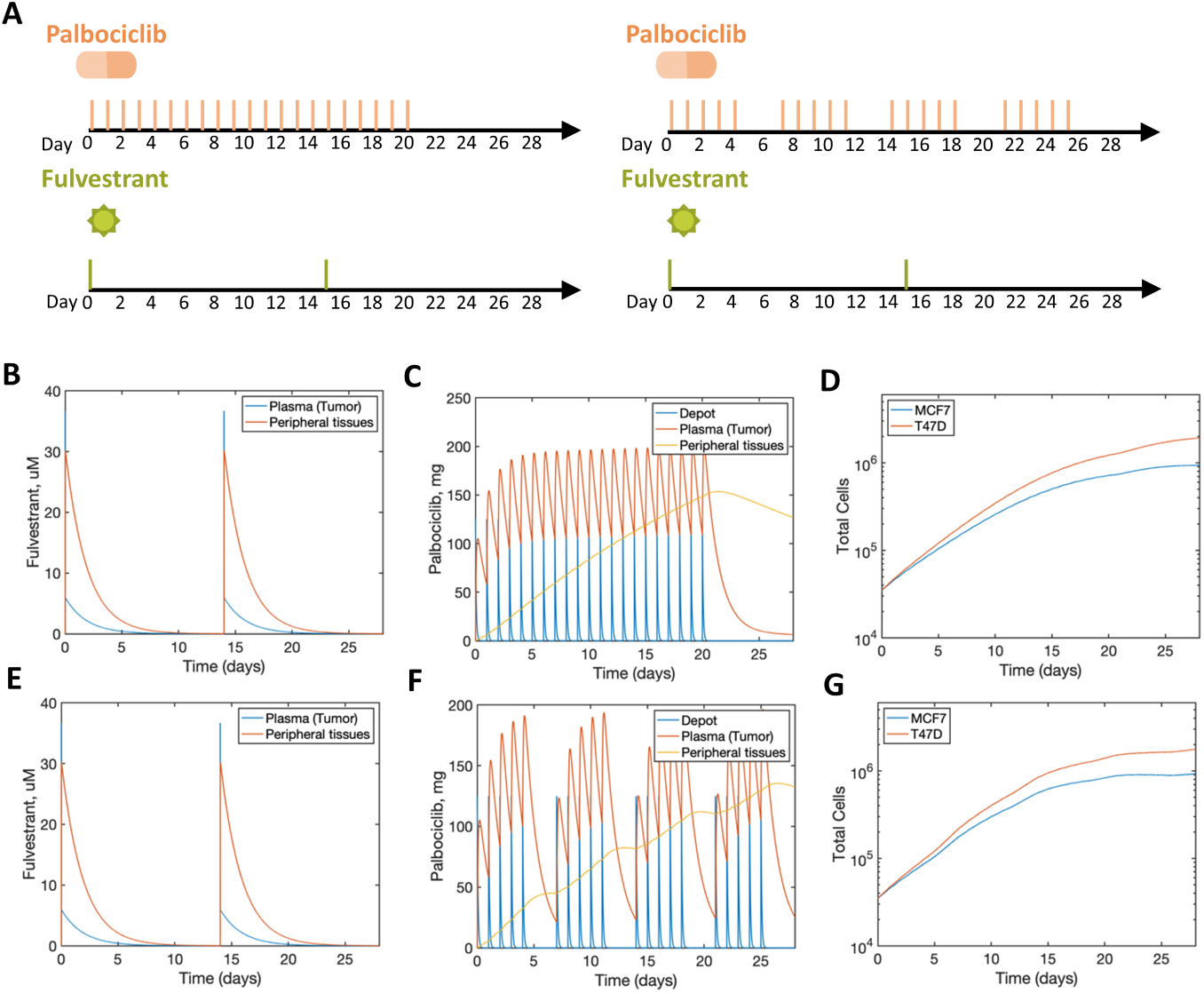
Comparison of alternate protocols for combination therapy. (A) Two established protocols are being considered for combination palbociclib and fulvestrant treatment denoted by this schematic. (B)-(D) Tumor growth dynamics with 21 days on of palbociclib with 7 days of rest. (B) and (E) Fulvestrant pharmacokinetic model (Eqs. 9-11). (C) and (F) Palbociclib pharmacokinetic model (Eqs. 6-8). (D) Tumor response to treatment. By comparing the total number of MCF7 and T47D cells at the end of treatment to the trial that did not receive treatment (Figure 14), we find a cell viability of 0.39 and 0.51, respectively. (E)-(F) Tumor growth dynamics with 5 days on of palbociclib with 2 days of rest, repeated for 28 days. (G) Tumor response to treatment. By comparing the total number of MCF7 and T47D cells at the end of treatment to the trial that did not receive treatment (Figure 15), we found a cell viability of 0.38 and 0.47, respectively. In Figure 16, we provide the corresponding effects function value over time.

### 3.2 Initial tumour composition has little impact on treatment outcomes

Given that our model predicted a slight reduction in T47D viability under shortened schedules for an average patient, we next interrogated how various levels of heterogeneity (e.g., intrinsic to the tumour population, PK, PD, and treatment scheduling) would affect outcomes. First, we explored the effect of the initial tumor composition and initial total cancer cell count on the outcomes of different treatment regimens (Figure 3 and Figure 17).

**Fig. 3.**
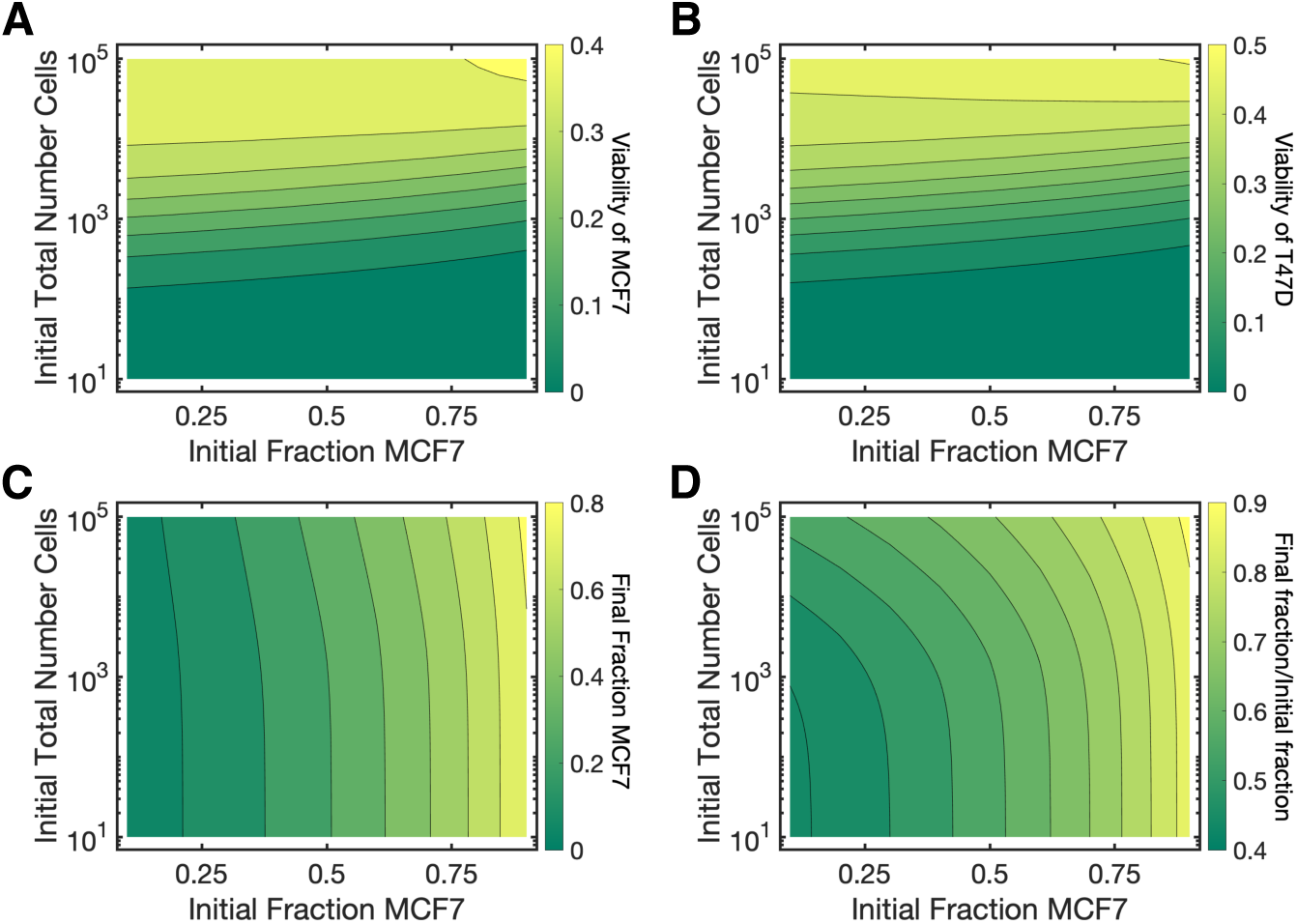
Results for varying initial tumor composition and total initial cell count, conventional treatment (i.e. 21 days on, 7 days off for palbociclib, Figure 2A). Initial fraction of MCF7 cell line (*ϕ*_*M*_) and total number of cells (*C*_*M*_ +*C*_*T*_) are varied over 0 < *ϕ*_*M*_ < 1 and 101 < *C*_*M*_ +*C*_*T*_ < 105. (A) Viability of the MCF7 line for conventional treatment over varied *ϕ*_*M*_ and *C*_*M*_ + *C*_*T*_ . Viability is calculated by comparing the total number of MCF7 cells with treatment compared to the total number of MCF7 cells without treatment after 28 days; both trials have the same initial conditions and only differ in whether treatment is administered. (B) Viability of the T47D line for conventional treatment over varied *ϕ*_*M*_ and *C*_*M*_ + *C*_*T*_ . Viability for T47D is larger than that of MCF7. (C) The final fraction of MCF7 cell line (*ϕ*_*M*_) after the 28 days of treatment. (D) The final fraction of MCF7 cell line (*ϕ*_*M*_) after the 28 days of treatment compared to the initial fraction.

To isolate the effect of the initial tumour composition, we set all model parameters in both the palbociclib and fulvestrant pharmacokinetic models and their pharmacodynamics to be their mean values (Table 1-3), as in the previous section. We then varied the initial fraction of MCF7 cells (*ϕ*_*M*_), i.e., the less aggressive cell type, from 0 to 1 and the total initial cell count (*C*_*M*_ + *C*_*T*_) from 10^1^ to 10^5^ cells. We sampled 440 parameter values with different *ϕ*_*M*_ and *C*_*M*_ + *C*_*T*_ within these ranges.

We found that the cell viability, defined as stated earlier by comparing the total cells at the end of treatment to the total number of cells under no treatment, and final fraction (*ϕ*_*M*_ after 28 days of treatment) over these varying initial conditions showed decreased T47D viability for the shortened treatment (i.e., shortened vs conventional, see Figure 3 and Figure 17). At lower initial cell counts, our model predicted that the more aggressive T47D cells were more likely to take over at the end of treatment (Figure 3). Our predictions show that as the initial number of cells increased, so too did the cell viability. This implies that with more cells, the drug combination becomes less effective. In both regimens, there appeared to be a switching point for the initial cell fraction above which MCF7 cells can take over (∼0.75). Though T47D viability did decrease with the shortened treatment, there was not an exceptional difference between the shortened and conventional treatment, indicating that the dosing schedules are more dependent on other factors of variability, i.e., pharmacokinetic and/or pharmacodynamic.

### 3.3 The effects of pharmacokinetic variability are determined uniquely through fulvestrant interindividual variability

To quantify the effects of interindividual variability in PK parameters on the outcomes of both MCF7 and T47D cells, we simulated the standard dosing regimen of each drug in the population of virtual patients defined by our estimated population PK (PopPK) models (see Figure 4, Figure 18).

**Fig. 4.**
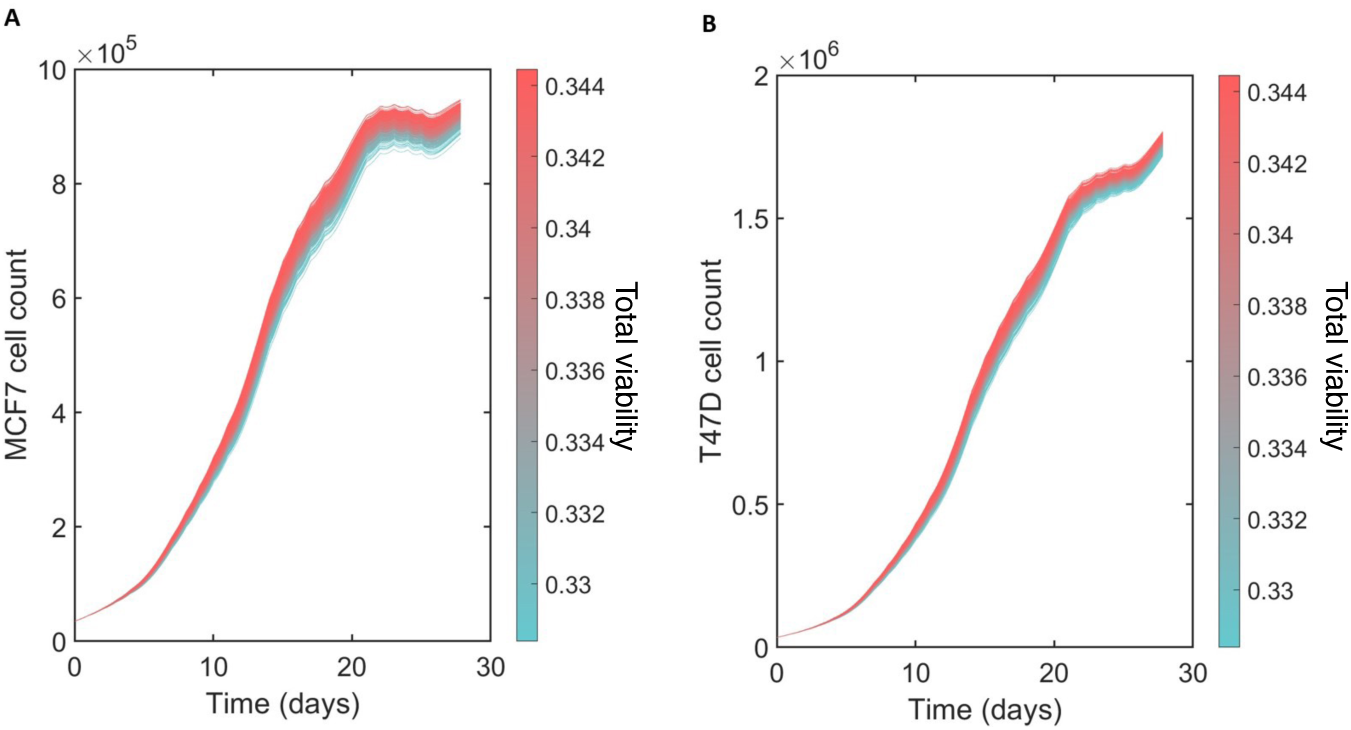
Predicted outcomes on standard regimen in palbociclib virtual patient cohort. (A) MCF7 and (B) T47D cell counts over the course of a standard treatment regimen with variation in palbociclib pharmacokinetic parameters summarized in Figure 18. Colour bar: viability of T47D cells.

For palbociclib, our results suggest that variability in the pharmacokinetic parameters has a negligible influence on tumour growth outcomes for both cell types (Figure 4 and 19). Interestingly, examining the palbociclib parameters by classifying patients as either responders or non-responders based on their predicted terminal T47D cell count, we see a clear distinction in the cohort’s value for *k*_*el*_, which is high for those with high terminal concentrations and low for those without (Figure 5).

**Fig. 5.**
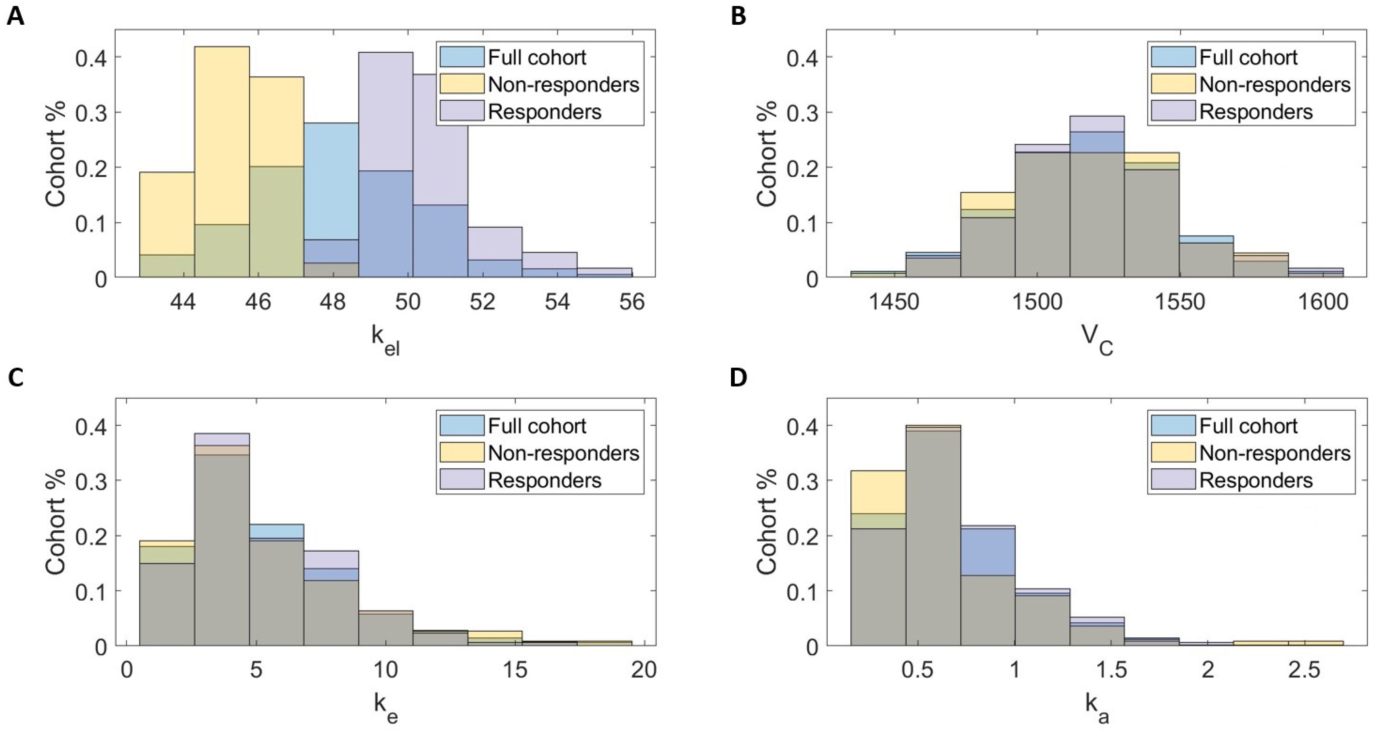
Some palbociclib PK parameters differ between responders and non-responders. Virtual patients were classified as responders or non-responders based on the predicted terminal T47D cell count of each virtual patient. Upon performing a two-sided Kolmogorov-Smirnov test for each parameter between these two subcohorts, we found significant differences in (A) the elimination rate (*k*_*el*_) and (D) the absorption rate (*k*_*a*_) and observed no significant differences in (B) the central volume (*V*_*C*_) and (C) the intercompartmental clearance rate (*k*_*e*_).

In contrast, our results suggest that fulvestrant pharmacokinetic variability has a significant impact on tumour growth outcomes for both cell types (Figure 6A and Figure 6B). Distribution of fulvestrant PK parameters in the virtual patient cohort are provided in Figure 20. We observed that virtual patients who sustained high concentrations of fulvestrant over the treatment period has significantly and consistently lower tumour growth as compared to virtual patients who more rapidly cleared the drug (Figure 6C).

**Fig. 6.**
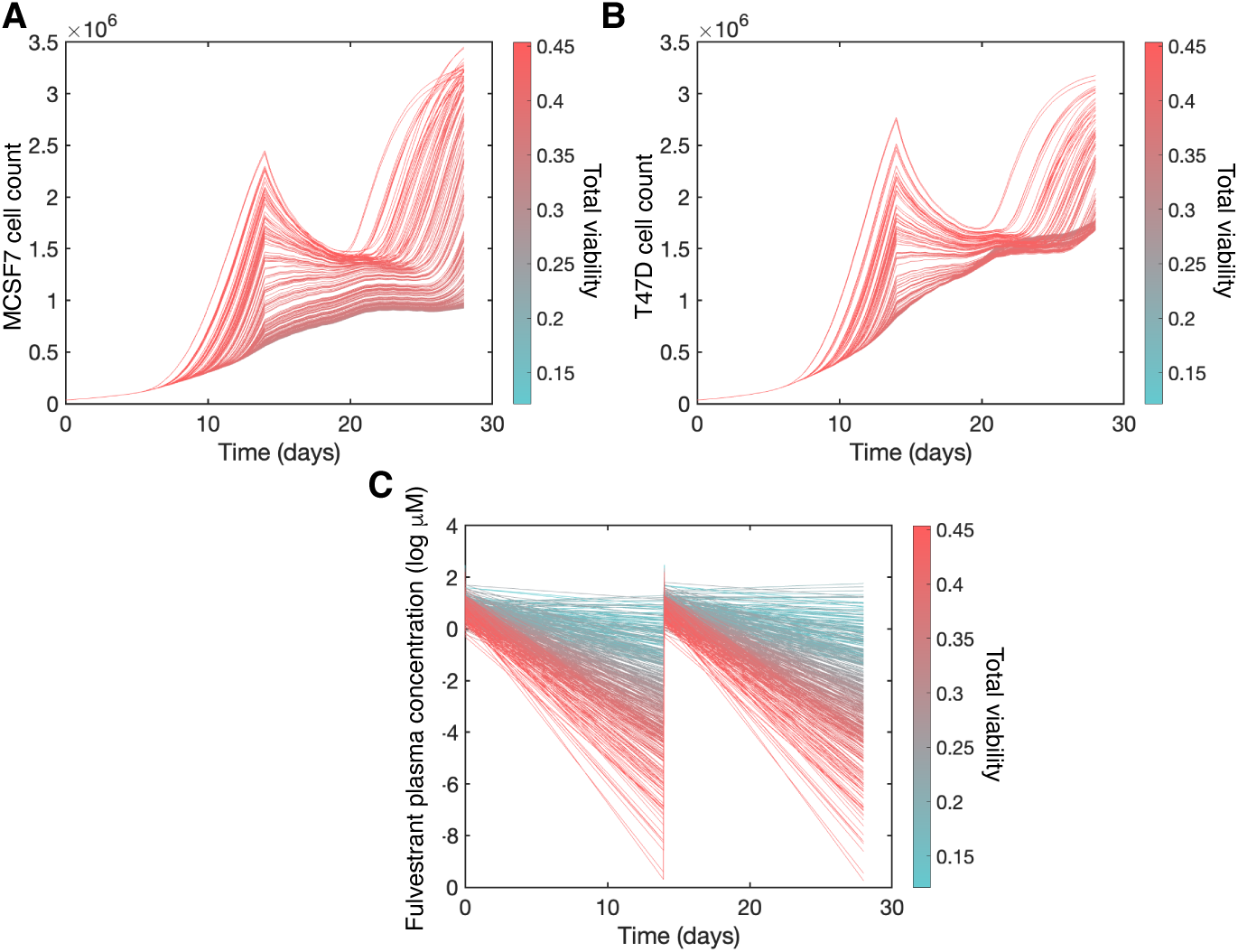
Spaghetti plots for fulvestrant virtual patients. Predicted dynamics for 438 patients in fulvestrant virtual patient cohort after treatment with 125 mg of fulvestrant on days 1 and 15. (A) MCF7 cells, (B) T47D cells, and (C) fulvestrant concentrations. In all, colour bar indicates total tumour viability.

Given the clear relationship between terminal fulvestrant concentrations and outcomes in our virtual patients, we defined virtual patients with “high concentration” to be those with terminal fulvestrant concentrations above -1.4 log(*μ*M) and those with “low concentration” as those with concentrations below -3.97 log (*μM*) (Figure 7). Using a two-sided Kolmogorov-Smirnoff test to test for statistically significant differences in distributions between these two subcohorts, we found significant differences in all fulvestrant PK parameters between these groups. This suggests that not only is fulvestrant the key driver of PK heterogeneity (as compared to palbociclib), but that differences in final tumour viability were related to higher *t*_*k*0_, *V*_*D*_, *k*_*el*_, and *k*_21_ values, and lower *k*_12_ values than those virtual patients who were predicted to have strong responses to fulvestrant treatments (Figure 7).

**Fig. 7.**
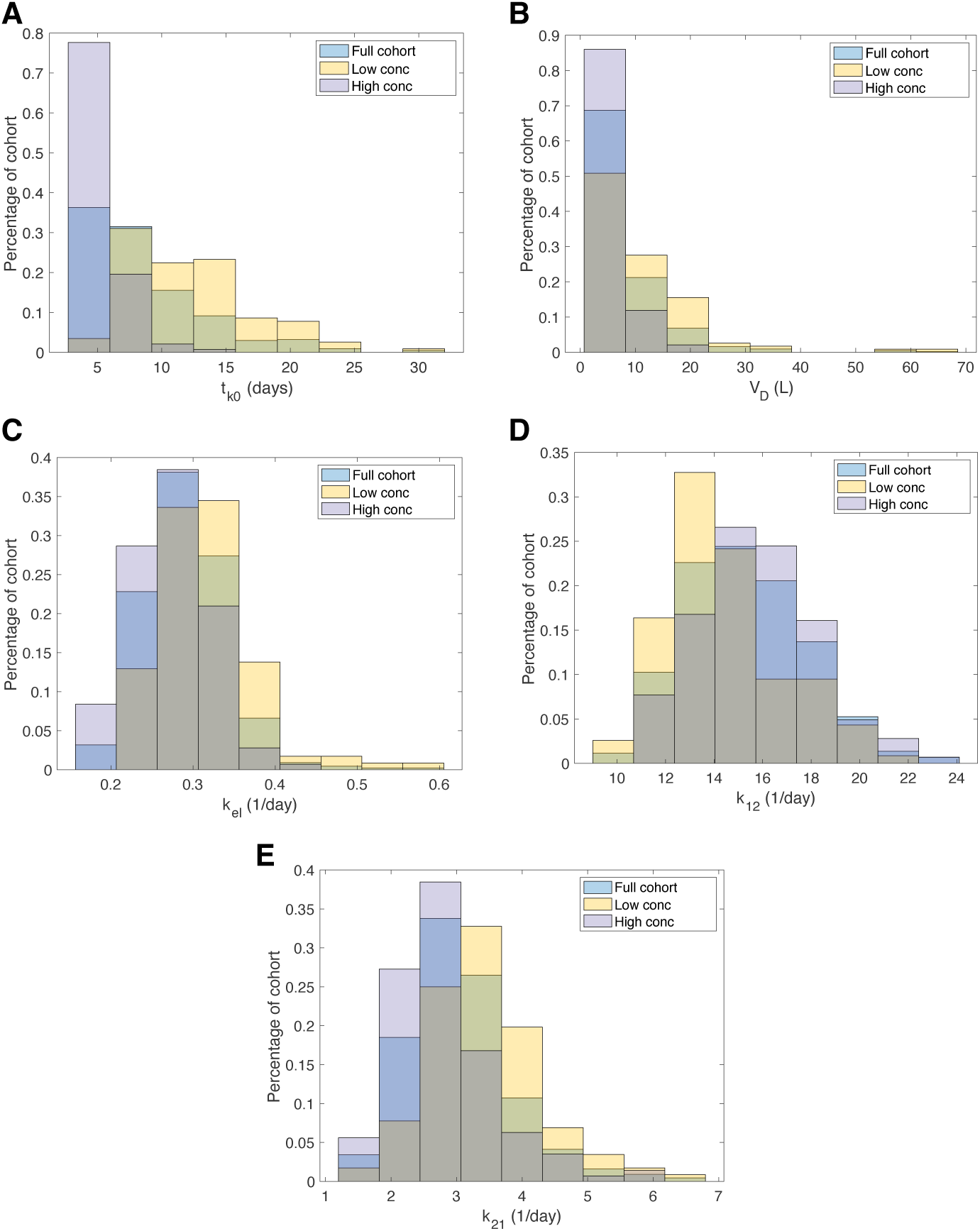
Fulvestrant PK parameters differ between virtual patients with high terminal fulvestrant concentrations and those with low terminal concentrations. We classified virtual patients as “high concentration” or “low concentration” based on the predicted terminal fulvestrant concentration of each virtual patient (Figure 6C). Using a Kolmogorov-Smirnoff test for differences in distributions, we found significant differences in the (A) absorption delay (*t*_*k*0_), (B) central volume of distribution (*V*_*D*_), (C) rate of elimination (*k*_*el*_), (D) rate of transfer from central to peripheral compartment (*k*_12_), and (E) rate of transfer from peripheral to central compartment (*k*_21_). In legends, “high conc” corresponds to those virtual patients with high terminal fulvestrant concentrations, and “low conc” to those with low terminal fulvestrant concentrations.

### 3.4 Variability in each drug’s maximal effect drives heterogeneity in outcomes

To next explore the effect of pharmacodynamic variability on treatment outcomes, we fixed all model parameters to be that of an average patient (see Table 1-3) except the pharmacodynamic parameters in Eq. 4-5 noted in *p̂*. We generated 400 parameter sets within this range. We then examined whether we could discern a relationship between each individual’s response to treatment and their inherent pharmacodynamic response. For each virtual patient, the standard treatment protocol was simulated, and the corresponding MCF7 and TD47 cell counts were recorded (Figure 8A) along with the total number of tumour cells (Figure 8B). We observed large variance in counts of both cell types across the cohort, but our model did not predict tumour eradication for any patient (Figure 21).

**Fig. 8.**
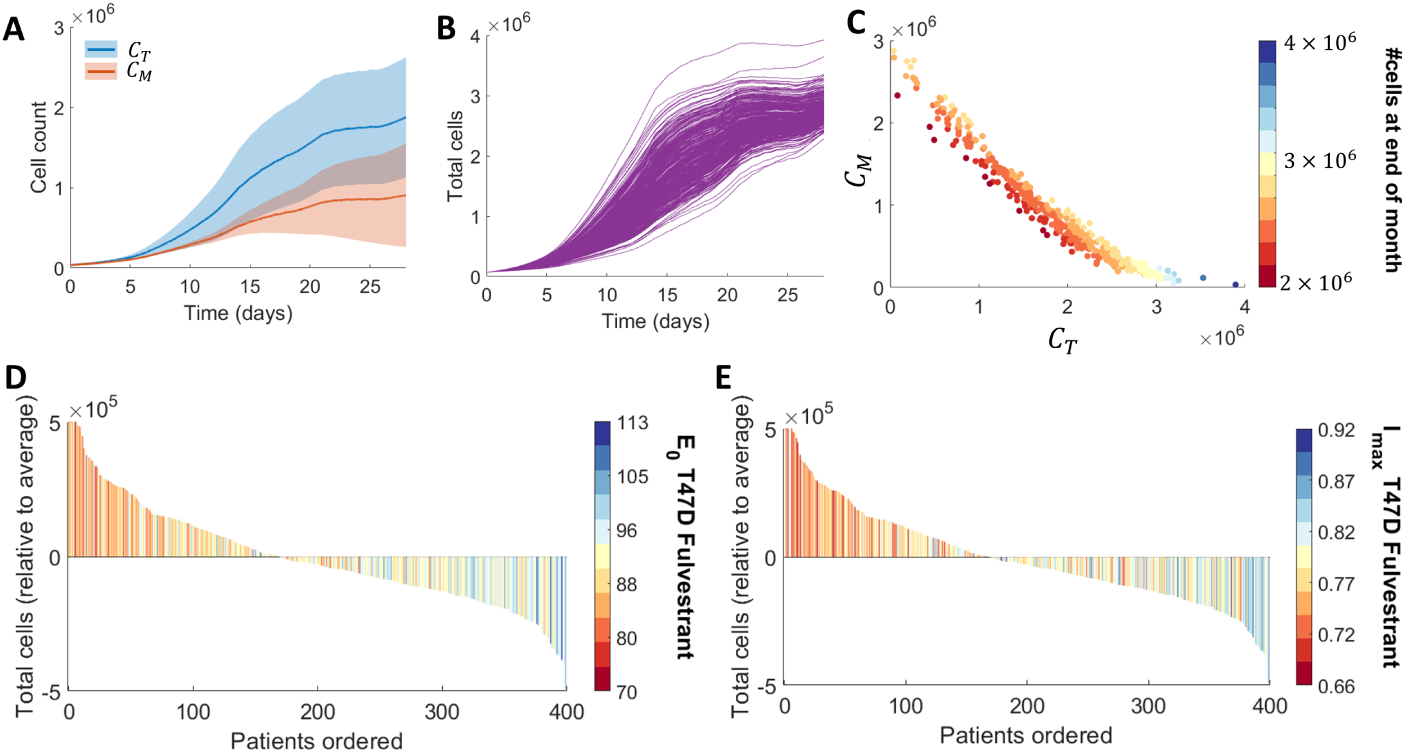
Virtual cohort investigation into the effect of pharmacodynamics on combination treatment. 400 virtual patients were generated with varying pharmacodynamic parameters (see *p̂*). A 5-day- on, 2-day-off paclbociclib regimen combined with two fulvestrant dosages was considered. (A) Cell counts for MCF7 and T47D cells over time plotted as mean and standard deviation of patient cohort. (B) Individual patient trajectories for total cell count *C*_*M*_ + *C*_*T*_ . (C) A scatter plot of the final number of each cell type after treatment, coloured by the corresponding total number of cells after treatment. (D)-(E) Waterfall plots for patient specific *E*_0,*T f*_ and *I*_*max*,*T f*_ against the total cells relative to the cohort average. Color bar corresponds to the value of each patient’s parameter normalised to a range between 0 and 1.

To examine the correlation between final number of MCF7 and T47D cells at the end of the treatment, we next plotted the total number of MCF7 and T47D cells at the end of treatment (Figure 8C) for each generated parameter set in our ensemble. Our results clearly show that as the final amount of MCF7 cells decreased, there was a corresponding increase in the final T47D cell count, and vice versa. In other words, final MCF7 and T47D were predicted to have an inverse linear relationship when pharmacodynamic variability was considered. We also correlated the final MCF7 and T47D cell counts with the final tumour size and found that the largest tumours are those that are predominately made up of T47D, whereas the smallest tumours are a mixture of both cell types.

We then examined which pharmacodynamic characteristics were the major drivers of final tumour size. We found that *E*_0_ and *I*_*max*_ for T47D (i.e. *E*_0,*T f*_,*I*_*max*,*T f*_) were most correlated with the final tumour size (Figure 8D-E), as we predicted that small values of either parameter would correspond to large tumour sizes relative to the average (Figure 22). All other parameters were not found to contribute significantly to the final tumour size.

### 3.5 Examining the long-term effect of variation on the combination protocol

Finally, with our understanding of the effects of cell-intrinsic, pharmacokinetic, and pharmacodynamic variability on combination palbociclib and fulvestrant therapy, we explored alternative treatment regimens to study whether we could improve upon the conventional and investigational schedules using virtual clinical trials of three different dosing regimens. On the current standard-of-care 3-weeks-on/1-week-off dosing schedule of palbociclib, numerous patients have been reported to develop grade 3 or higher degree of neutropenia [13]. This adverse event could result in dose reduction or treatment discontinuation [13]. Further, it has been hypothesized that the one week off-drug in the conventional combination schedule could potentially lead to an increase in the retinoblastoma tumor suppressor protein (Rb) [13].

We therefore explored alternative schedules with the aim of reducing the time off-drug (as compared to the conventional regimen), with limited dose intensity, to minimize off-target effects (Regimen 1). Additionally, clinical reports suggest that fulvestrant is mostly likely to cause acute liver injury [6, 27]. To reduce the risk of hepatotoxicity, we virtually reduce the dose level of fulvestrant and test the effectiveness of this dual-agent combination therapy in Regimens 2 and 3. Thus, we compared the following three schedules:

**Regimen 1**: 125 mg of oral palbociclib administered once daily for 5 consecutive days, followed by 2 days off, plus 500 mg of intramuscular (IM) fulvestrant administered every 14 days for the first 3 injections and then every 28 days.

**Regimen 2**: 125 mg of oral palbociclib administered once daily for 5 consecutive days, followed by 2 days off, plus 250 mg IM of fulvestrant administered every 7 days for the first 5 injections and then every 14 days.

**Regimen 3**: 500 mg of oral palbociclib administered once daily for 5 consecutive days, followed by 2 days off, plus 250 mg IM of fulvestrant administered every 7 days for the first 5 injections and then every 14 days.

Simulations of Regimen 1 suggest that this schedule leads to the competitive exclusion of aggressive T47D cells. Selective killing of the therapy sensitive cells removes competitive restriction of MCF7 cells (Figure 9A). The troughs of the fluctuating total cell loads come down to 3×10^7^, while the peaks still reach a high level (Figure 9B). In contrast, by reducing the dose of fulvestrant, Regimen 2 did not lead to competitive exclusion of the T47D cells (Figure 9C), but resulted in an overall significant decrease in the total number tumour cells (Figure 9D). We further found that Regimen 3 did not lead to competitive exclusion of the T47D cells (Figure 9E), and our model predictions suggest that the total number of cancer cells from both lines would continue to decrease (Figure 9F).

**Fig. 9.**
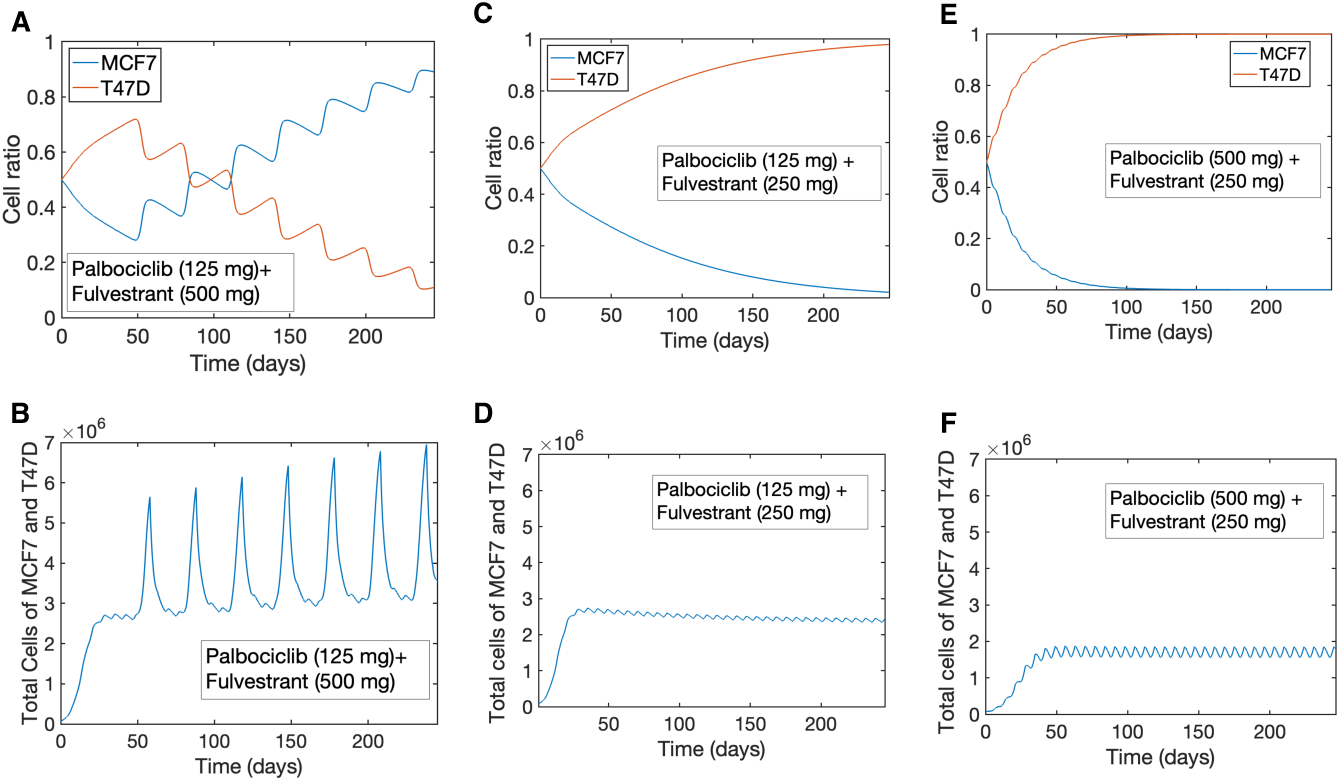
Investigating long-term dynamics of Regimens 1, 2, and 3. (A), (C), and (E) cancer cell ratios after treatment with Regimens 1, 2, and 3. Blue lines: ratio of cancer cell line MCF7 given by *C*_*M*_/(*C*_*M*_ + *C*_*T*_); red lines: ratio of cancer cell T47D given by *C*_*T*_ /(*C*_*M*_ + *C*_*T*_). (B), (D), and (F) comparison of the total cell load of both cell lines after treatment with the three regimens. B) Regimen 1; D) Regimen 2; F) Regimen 3.

Overall, we found that increasing the dose level of palbociclib within acceptable toxicity levels could achieve a lower level of total cancer cell load. Importantly, based on our simulations of our 3 dosing regimens, it is possible that higher doses/concentrations of fulvestrant could cause competitive exclusion of the T47D cell line. As a result, the relative strength of the less-aggressive MCF7 cells may in fact inhibit the efficacy of the palbociclib-fulvestrant combination therapy.

## 4 Discussion

Heterogeneity is a key factor in cancer therapeutic planning, particularly when considering combination therapies that may have overlapping and interacting factors driving treatment responses. The interest in establishing different treatment regimens for palbociclib plus fulvestrant for the treatment of hormone-sensitive breast cancers gives rise to a number of questions relating to optimal scheduling. These include the various scales of heterogeneity and their impact on combination palbociclib and fulvestrant, i.e., cell-intrinsic, pharmacokinetic, and pharmacodynamic. Understanding the contributions of each of these elements to tumour responses helps to establish new, and perhaps more potent and less toxic, therapeutic regimens. In this work, we used a simple model of interacting cells to quantify these contributions to help guide preclinical studies of palbociclib plus fulvestrant.

Considering a tumour composed of lesser and more aggressive cells (i.e., MCF7 and T47D cell lines), each type sensitive to a different degree to each drug, we predicted the overall tumour cell population and composition after treatment under variable initial fractions. Our predictions showed that the initial cell fractions have little impact on the final tumour composition after treatment on either the shortened (i.e., 5 days on of palbociclib with 2 days of rest, repeated for 28 days) or conventional 21 days on of palbociclib with 7 days of rest schedule. This is encouraging, in the sense that it suggests that it is primarily PK/PD variability controlling outcomes and these can be more easily modulated to provide better results.

When considering both PK and PD heterogeneity through the generation of virtual patients, we found that palbociclib PK variability alone had little impact on outcomes, whereas the PKs of fulvestrant (as a cytotoxic agent) was strongly determinate of final tumour compositions. This is perhaps expected, as palbociclib acts to freeze the cell cycle rather than induce apoptosis. Our results further show that palbociclib and fulvestrant are truly synergistic when given in combination, with each being less effective on its own. Lastly, we used our investigations of the impact of various scales of heterogeneity to propose three alternative regimens to conventional and shortened. These regimens were designed to account for the undesired side-effects of each drug through dose fractionation. Our model predictions suggest that it is possible that fulvestrant could cause competitive exclusion of the MCF7 (or less aggressive) cells composing the tumours in our study. Indeed, our results showed that the more aggressive T47D cells act to inhibit the efficacy of the palbociclib-fulvestrant combination therapy, acting similarly to drug tolerant cells despite us not considering resistance in our study. Moreover, within acceptable toxicity levels, increasing the dose level of palbociclib could achieve better outcomes with respect to final tumour size.

In our model, we implemented a logistic growth function to model tumour growth. While Gompertzian tumour growth returned a similar Akaike information criterion (AIC) and was able to capture the data (Figure 11), we do not anticipate a large difference in our predictions between the two growth models. This is largely due to the fact both exhibit sigmodal style growth to a carrying capacity.

**Fig. 10.**
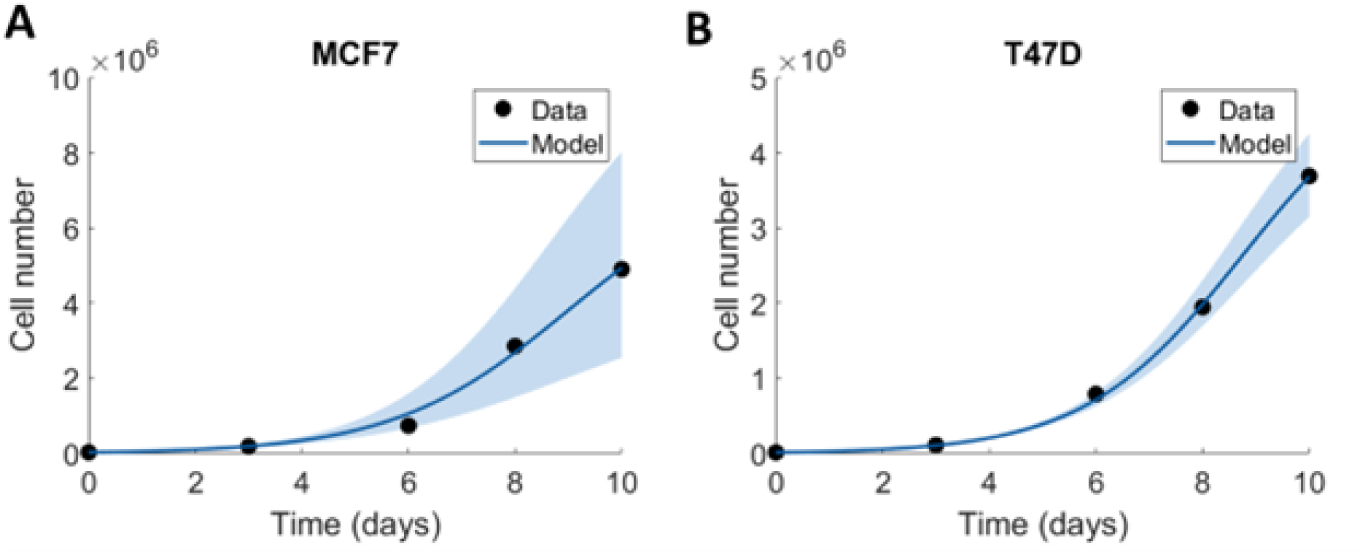
Fit of logistic growth to cell count measurements for MCF7 and T47D from Vijayaraghavan et al [35]. A logistic growth curve was fit to the cell count measurements to obtain a cell growth rate *r*_*i*_ (MCF7 and T47D) and a cell carrying capacity *K*_*i*_ (MCF7 and T47D). The fit is given as a solid line with a shaded 95% confidence interval. The data is represented as solid points. The resulting parameter values for the fits are given in Table 3.

**Fig. 11.**
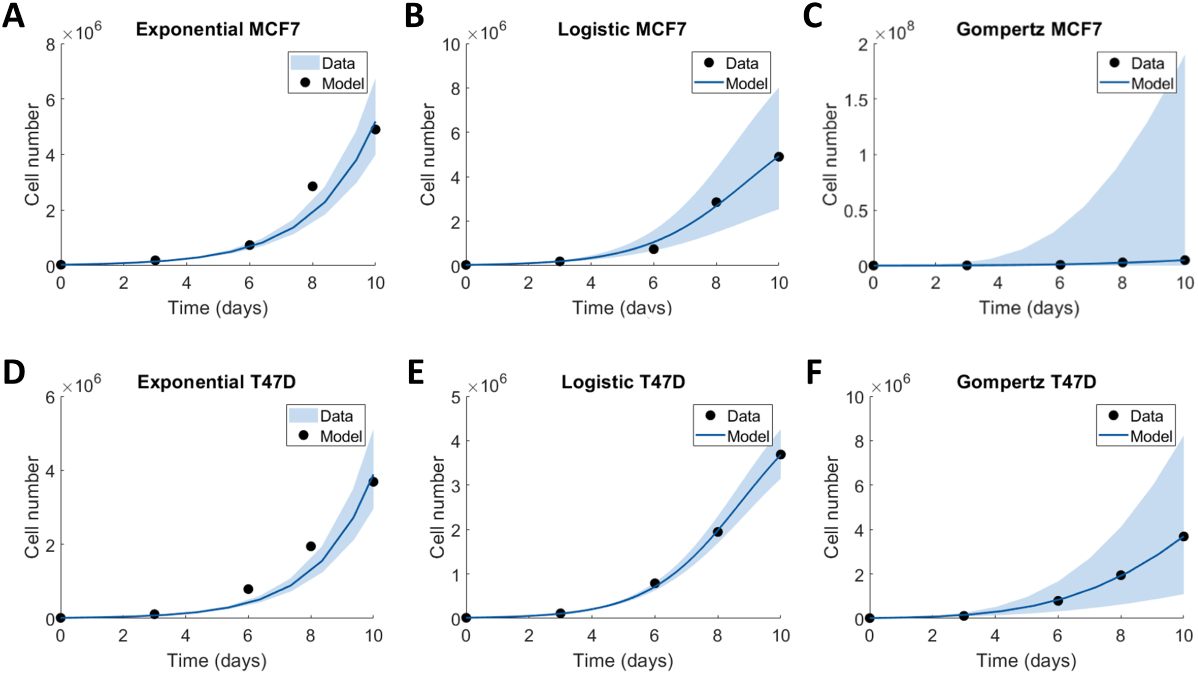
Comparative analysis of model selection to MCF7 and T47D cell count data. We compared the least-squares fit for (A,D) exponential growth, (B,E) logistic growth and (C,F) Gompertzian growth. We calculated the corrected Akaike Information Criterion (AIC) for each figure which returned: (A) 134.3, (B) 131.1, (C) 134.5, (D) 131.7, (E) 116.5, and (F) 115.0. We also considered the confidence intervals plotted for each model fit. Given that the Gompertzian growth has wider confidence intervals compared to logistic growth, we concluded that logistic growth was a good model choice for tumour growth.

**Fig. 12.**
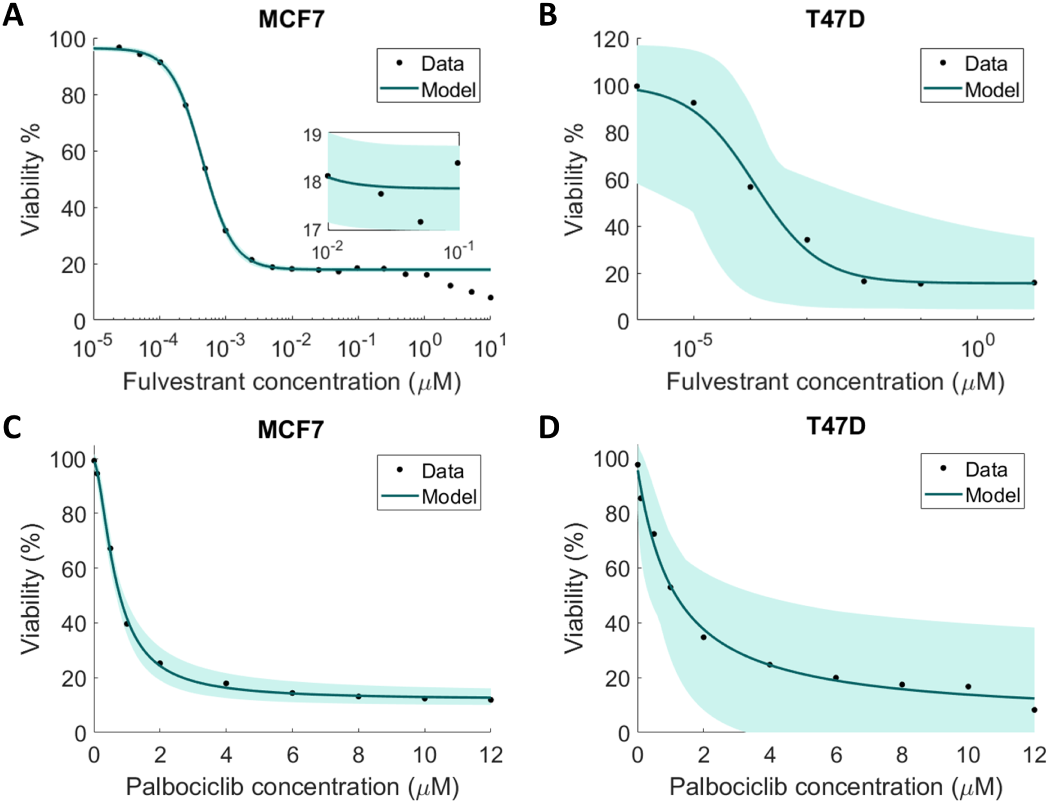
Fit of drug effect parameters to cell viability measurements for fulvestrant and palbociclib on MCF7 and T47D. Cell viability measurements for (A) Fulvestrant on MCF7 cells [22] and (B) Fulvestrant on T47D cells [15]. Cell viability measurements for (C) palbocicblib on MCF7 and (B) T47D by Vijayaraghavan et al. [35]. The resulting parameter fits are in Table 3. In (A) the inset zooms in on the confidence intervals surrounding the data fit.

**Fig. 13.**
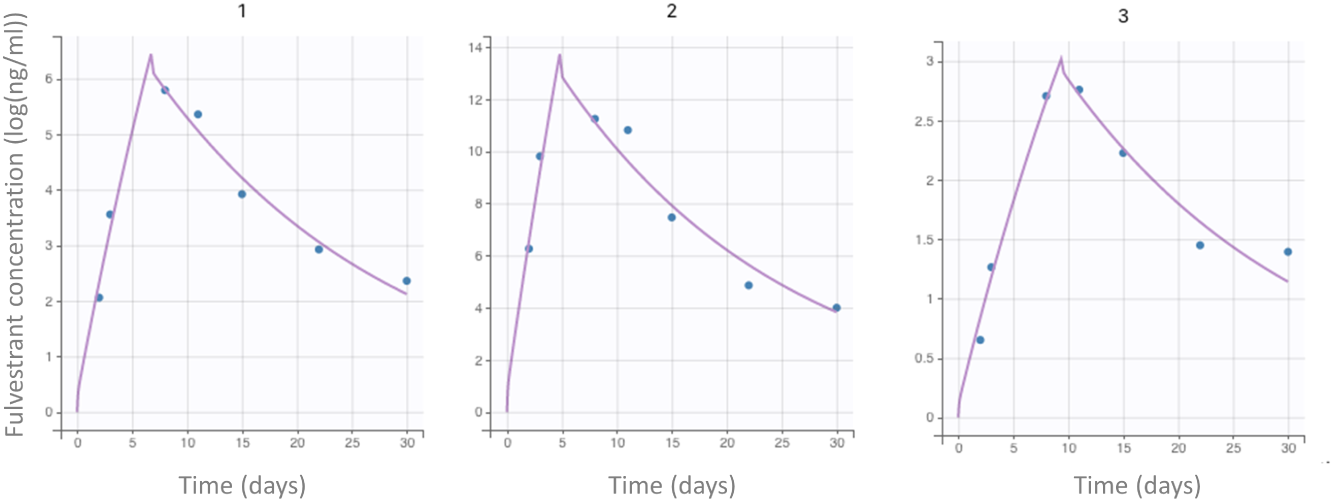
Fulvestrant pharmacokinetic parameter estimation fits. Population pharmacokinetic data from Robertson et al. [26] was pooled to extract the mean, and lower and upper bounds of the data. We then estimated PK parameters from Eqs. 6-8 in the Main Text to these data assuming lognormal distributions on parameters subject to IIV using a standard nonlinear mixed effects model in Monolix[16].

**Fig. 14.**
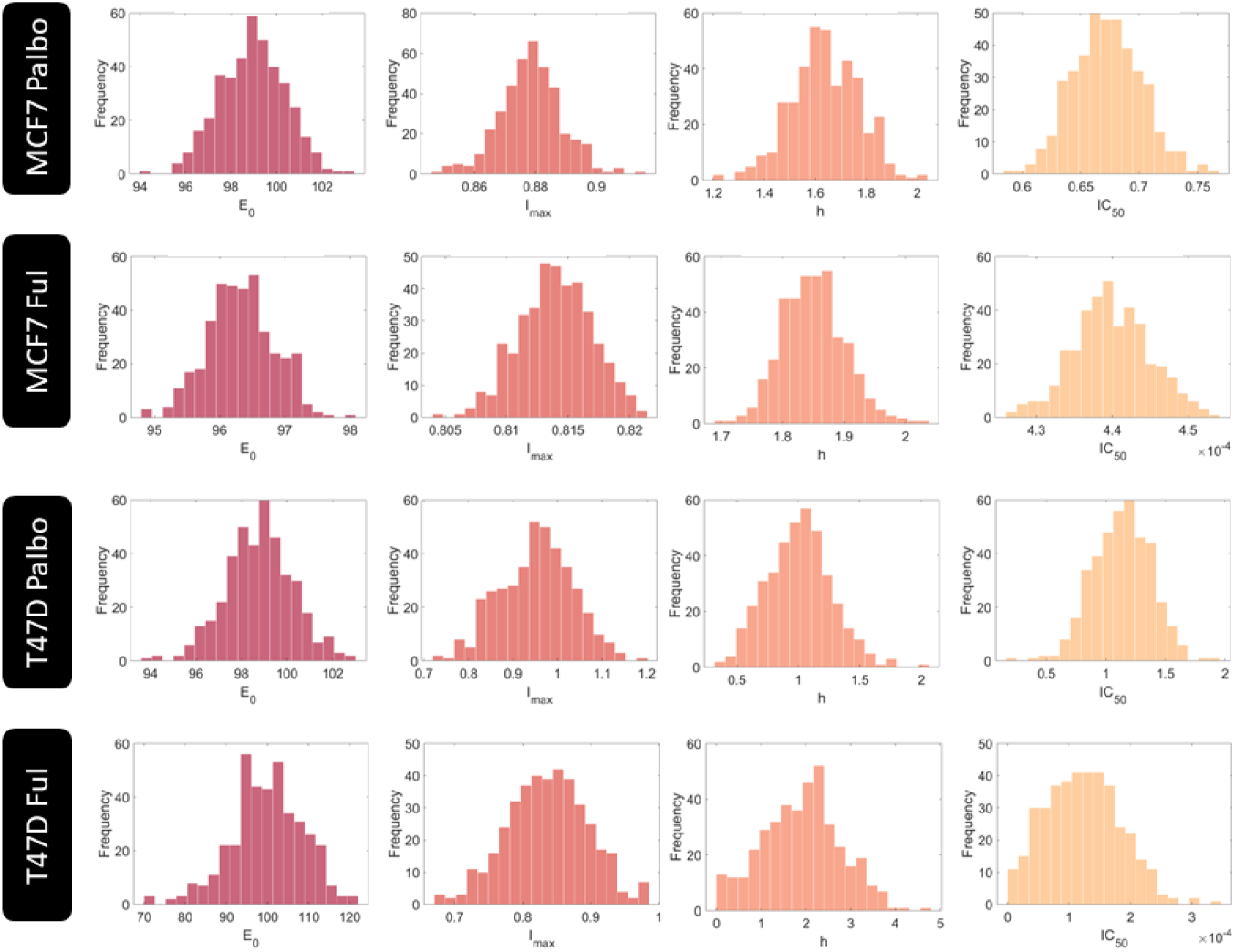
Virtual patient parameter values for investigation varying the drug effect parameters. Parameters relating the effect of fulvestrant and palbociclib on MCF7 and T47D were sampling from normal distributions as described in the methods to obtain 400 unique parameter combinations corresponding to 400 virtual patients.

**Fig. 15.**
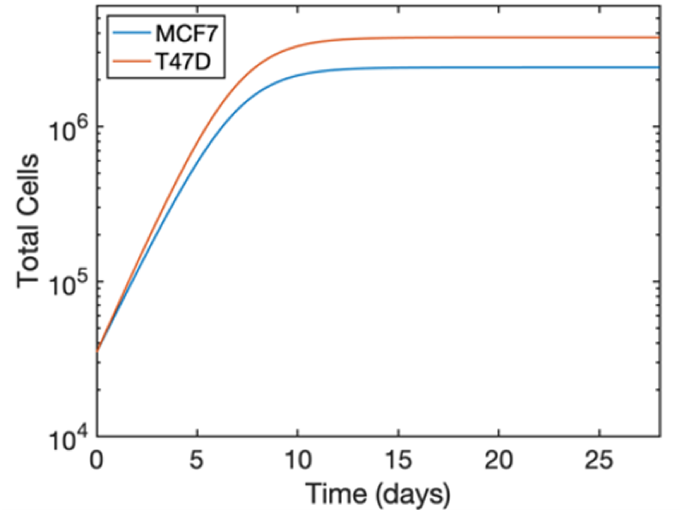
Tumor growth dynamics under no treatment. The total number of MCF7 and T47D cells, *C*_*M*_ and *C*_*T*_ respectively, were simulated in the absence of palbociclib and fulvestrant, i.e., *F* = *P* = 0.

**Fig. 16.**
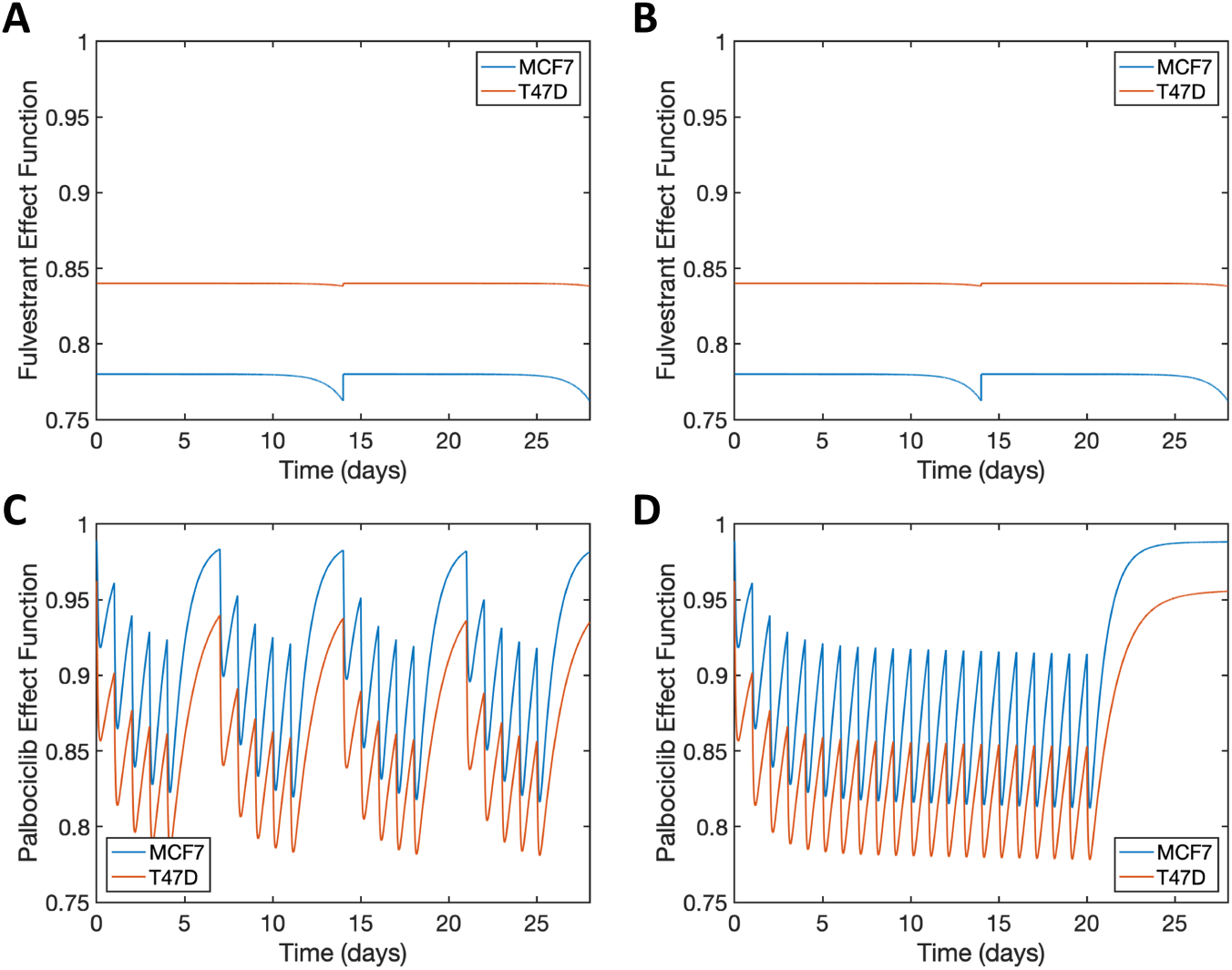
The effect function corresponding to the simulation in Figure 2. The effect function Eq. 1 for fulvestrant and palbociclib for the two dosage protocols considered in Figure 2: (A), (C) palbociclib 5 days on and 2 days off and (B), (D) palbociclib 21 days on 7 days off.

**Fig. 17.**
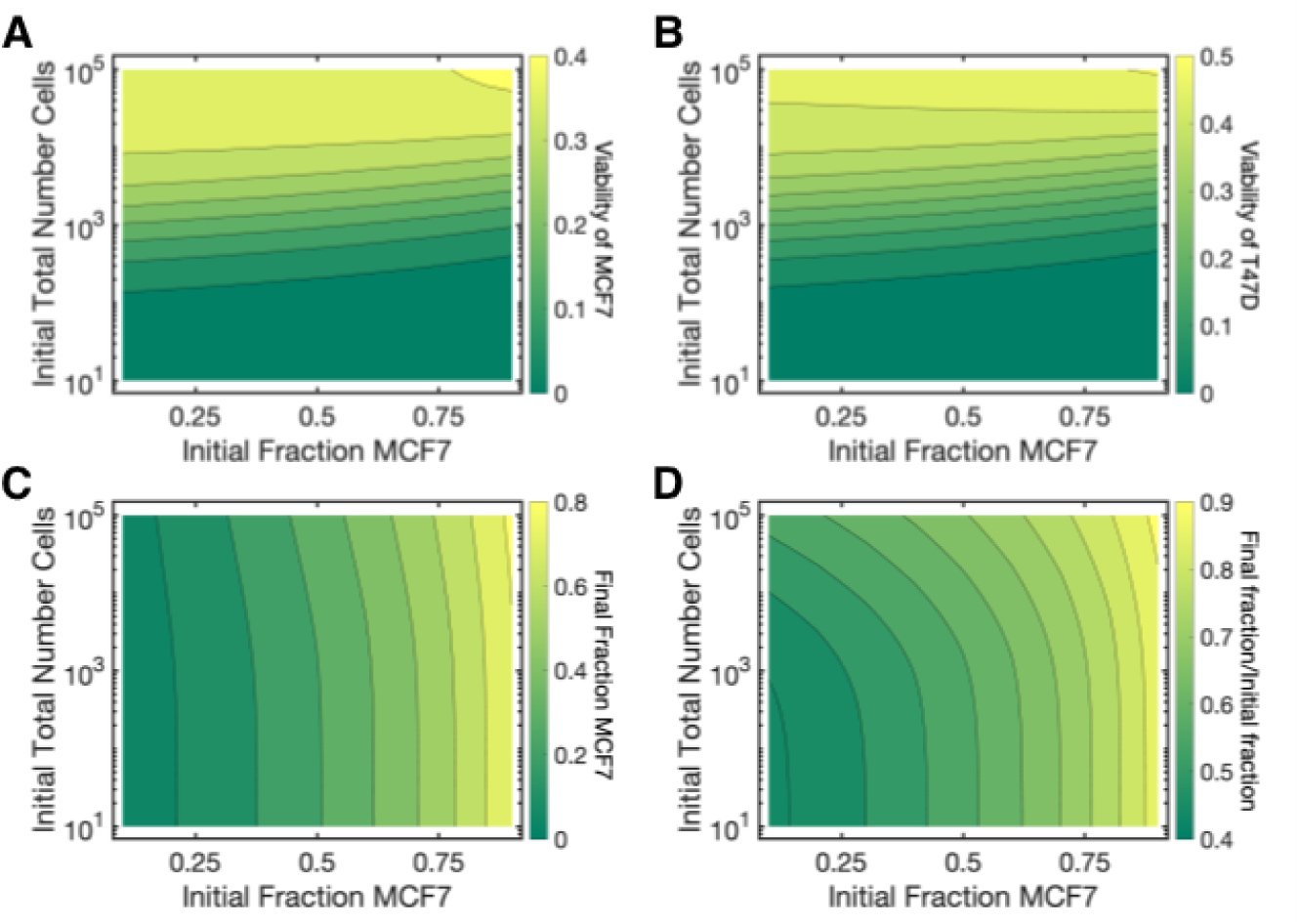
Results for varying initial tumor composition and total initial cell count, shortened treatment (i.e. 5 days on and 2 days off for palbociclib, Figure 2A). Initial fraction of MCF7 cell line (*ϕ*_*M*_) and total number of cells (*C*_*M*_ + *C*_*T*_) are varied over 0 < *ϕ*_*M*_ < 1 and 101 < *C*_*M*_ + *C*_*T*_ < 105. (A) Viability of the MCF7 line for shortened treatment over varied *ϕ*_*M*_ and *C*_*M*_ + *C*_*T*_ . Viability is calculated by comparing the total number of MCF7 cells with treatment compared to the total number of MCF7 cells without treatment after 28 days; both trials have the same initial conditions and only differ in whether treatment is administered. The overall viability of MCF7 is less for the shortened treatment, compared to the conventional treatment. (B) Viability of the T47D line for shortened treatment over varied *ϕ*_*M*_ and *C*_*M*_ + *C*_*T*_ . The overall viability of T47D is less for the shortened treatment, compared to the conventional treatment. (C) The final fraction of MCF7 cell line (*ϕ*_*M*_) after the 28 days of treatment. (D) The final fraction of MCF7 cell line (*ϕ*_*M*_) after the 28 days of treatment compared to the initial fraction. At lower initial total cell numbers, T47D has a greater propensity to overtake the cancer tumor and take up a greater fraction of the total tumor.

**Fig. 18.**
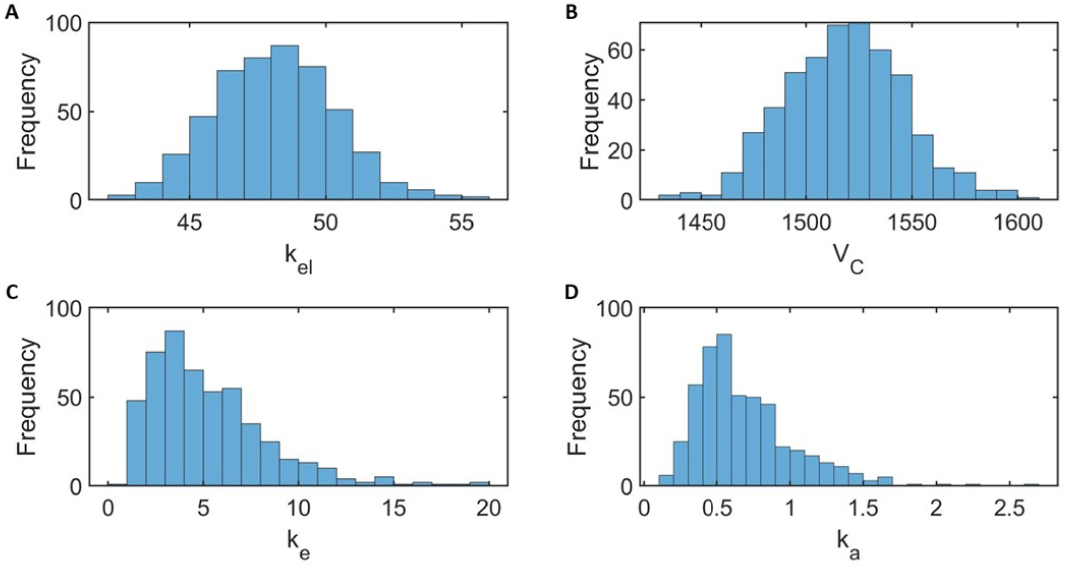
Distributions of palbociclib virtual patient parameters. Pharmacokinetic parameter distributions for (A) the elimination rate, (B) the central volume, (C) the intercompartmental clearance rate, and (D) the absorption rate of palbociclib constructed from 500 virtual patients sampled as described in the Methods. Experimental data suggests negligible variation in the parameter VP. This parameter is thus taken to be constant, hence the lack of distribution. The virtual patient population randomly generated to produce these distributions is maintained in the generation of Figure 5 in the Main Text.

**Fig. 19.**
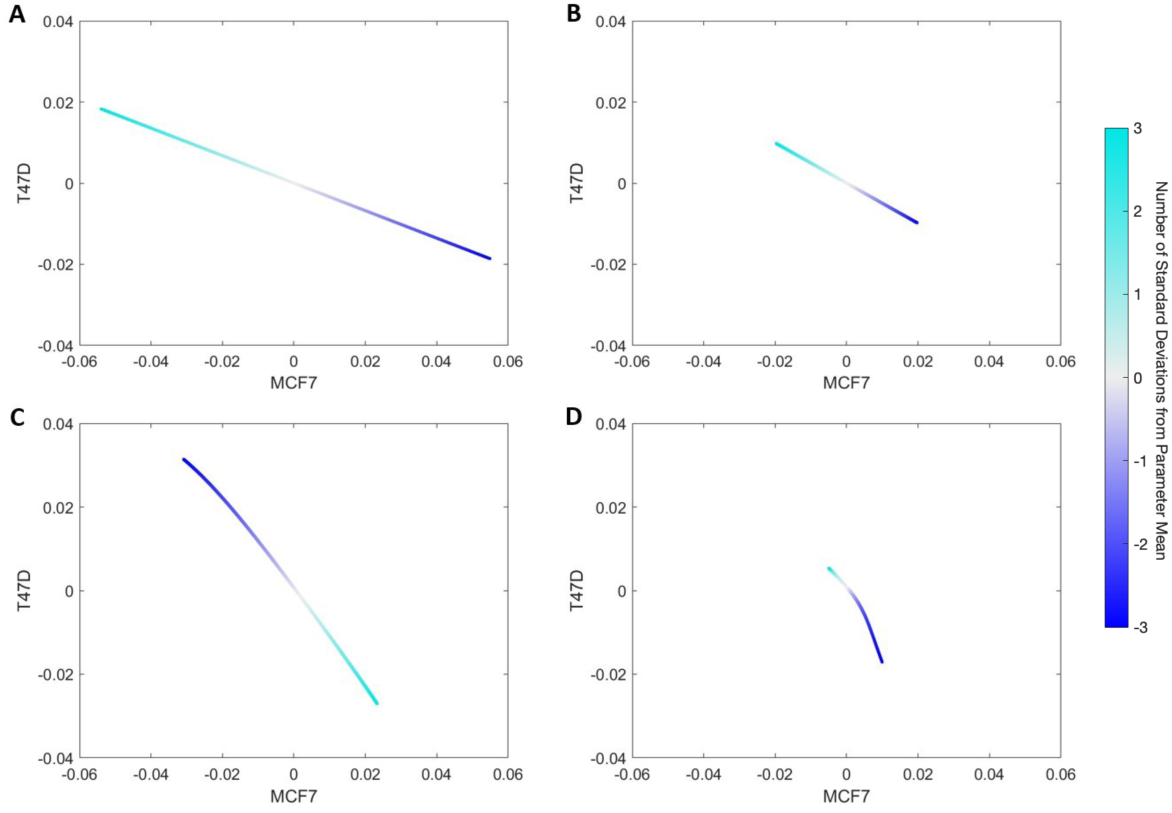
Palbociclib pharmacokinetic model sensitivity analysis. Parameter sensitivity of the palbociclib pharmacokinetic parameters: (A) *k*_*e*_*l*, (B) *V*_*C*_, (C) *k*_*e*_, and (D) *k*_*a*_. Curves convey changes in cell counts resulting from varying each parameter uniformly within three standard deviations of its mean according (Table 1). Numerical values on the axes correspond to the fractional deviation of an end-of-treatment cell count from its mean scaled by the fractional deviation of one standard deviation of a parameter from that parameter mean. Qualitatively, the distribution of a parameter non-negligibly influences final cell count if the curves above have coordinate values far from zero and close to one. Thus, our results show that all the parameter distributions have a negligible influence on the final cell counts.

**Fig. 20.**
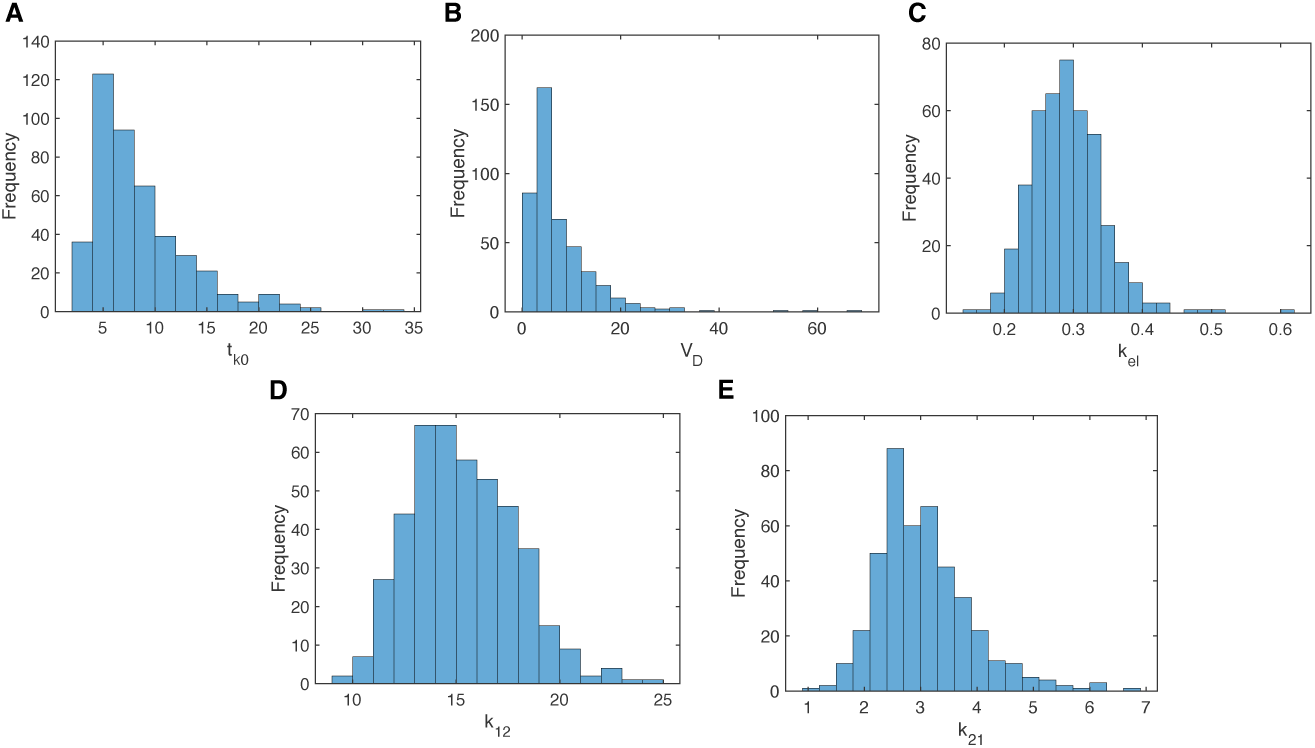
Distributions of fulvestrant virtual patient parameters. Pharmacokinetic parameter distributions for fulvestrant parameters (A) absorption delay (*t*_*k*0_), (B) central volume of distribution (*V*_*D*_), (C) rate of elimination (*k*_*el*_), and (D)-(E) rates of transit between central to peripheral compartments (*k*_12_ and *k*_21_). Distributions describe the 438 virtual patients in the fulvestrant virtual patient cohort, generated as described in the Methods (Main Text).

**Fig. 21.**
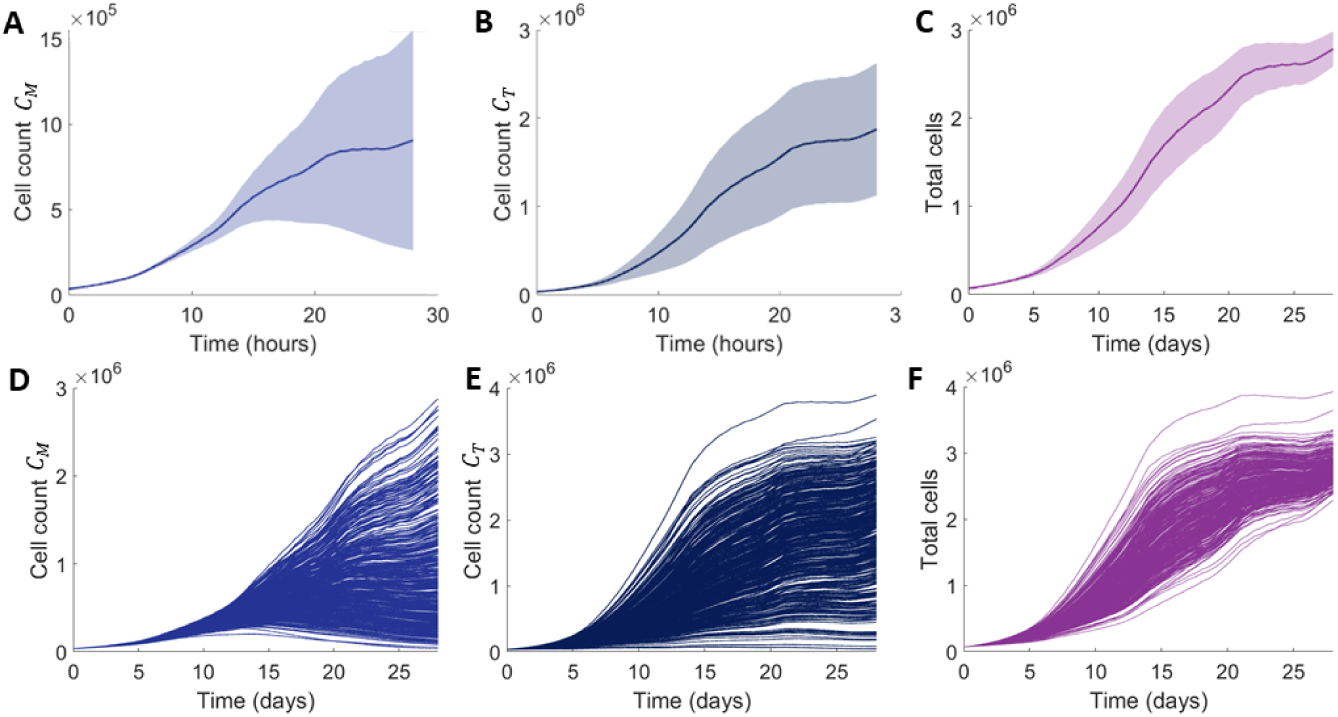
Results of simulating the virtual cohort with varying pharmacodynamics. The first show rows the mean and standard deviation for the 400 virtual patient cohort simulated with varying effect parameters (Figure 12). The second row is the corresponding individual patient trajectories.

**Fig. 22.**
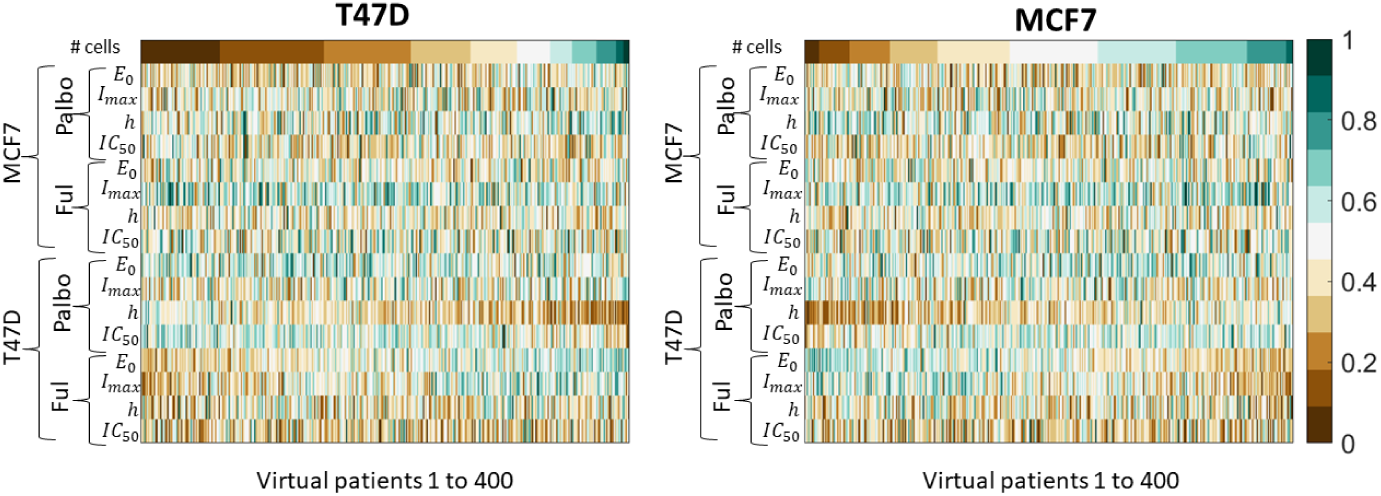
esults of simulating the virtual cohort with varying pharmacodynamics. Each column corresponds to a virtual patient where the patients are ordered by the number of cells (left: T47D and right: MCF7) at the end of the treatment. The rows correspond to the parameter value where the column is the normalised value for that parameter. The colour bar corresponds to the value of the parameter normalised between 0 and 1.

There are limitations to our approach.The lack of robust clinical data measuring this combination therapy presents a limitation in the reliability of our predictions. Given this is an exploratory study focused primarily on the effects of different sources of heterogeneity, we believe our results support further experiments into combination therapy that may be used in the future to validate our model predictions. Our parameterisation could be further validated with *in vivo* experiments. Future work could also look to refine the number of parameters in the model and introduce simpler terms to model the effect of the drugs on the cancer cell population or simplify the cancer growth function.

Though we considered heterogeneity in tumour composition, we did not include the mammary stem cell cascade [14, 30], nor does our model include the actions of the immune system. We opted for the simpler model studied herein to provide a straight-forward initial (more *in vitro*-focused) conceptualization of the impact of the many scales of heterogeneity affecting treatment outcomes under combination palbociclib and fulvestrant; future studies will include key *in vivo* factors impacting therapies. As seen in Fig. S3A, there is a loss of fidelity in the MCF7 fits for high fulvestrant concentrations. This is a byproduct of the standard nonlinear effects function (Eq. 1) used herein. As can be seen in the data, after an initial plateau beginning around 10^−2^ *μM*, there is a second dip in the observed viability around 1*μM*. Unfortunately, our effects function is unable to capture this second decrease, resulting in a higher predicted maximal effect than what is suggested in the data. To confirm the asymptotic behaviour of MCF7 cells under treatment with fulvestrant, it would be necessary to have more viability data above 1*μM*. Nonetheless, as our maximal effect is potentially slightly higher than in the data, our predictions are more conservative with respect to the overall treatment response to fulvestrant. We also did not consider the ways in which each degree of heterogeneity interacts with one another, opting instead to study each individually. This can be incorporated in subsequent iterations of our work.

Our study provides a roadmap for the continued study of CDK4/6 inhibitors and combination therapies in anti-cancer treatments more broadly. Despite using a simple model of tumour growth, our model’s predictions showed important perhaps unexpected behaviours, including how competition between less and more aggressive cells in a heterogeneous tumour impacts treatment scheduling. Ultimately, this work demonstrates the importance of merging mathematical modelling within preclinical studies to improve drug development considerations.

## Acknowledgements

The authors would like to thank the organizers, sponsors, and participants of the Collaborative Workshop for Women in Mathematical Biology: Mathematical Approaches to Support Women’s Health 2022, particularly the Institute for Mathematics and its Applications and UnitedHealth Group-Minnetonka, for bringing them together and providing support.

## Supplementary Information

### Supplementary analysis of the two cell-line model dynamics

To understand the dynamics of the system of ODEs used in our study, we carried out a linear steady state analysis to determine long-term stability of a simplified version of the system. We took the full model in Eq. 2-3 and considered only the effects on MCF7 and T47D cells of palbociclib:

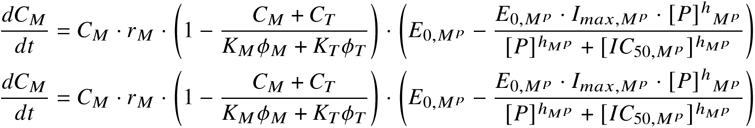

where

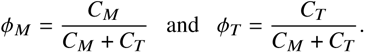

We assumed both cell lines share the same carrying capacity due to them co- existing in the same spatial location and hence feeling the same spatial limitations, that is *K* = *K*_*M*_ = *K*_*T*_ . For convenience for this analysis, we also assume that all pharmacodynamic parameters are equivalent for the two cell types, i.e., *I*_50,*M* *p*_ = *I*_50,*T* *p*_ = *IC*_50_, and ℎ_*M* *p*_ = ℎ_*T* *p*_ = ℎ, and *I*_*max*,*M* *p*_ = *I*_*max*,*T* *p*_ = *I*_*max*_, and *E*_0_ = *E*_0,*M* *p*_ = *E*_0,*T* *p*_, etc. For convenience, we denote *r*_*M*_ *E*_0_ = *r̃* _*M*_ *andr*_*T*_ *E*_0_ = *r̃* _*T*_ .

The model has three isolated equilibrium points

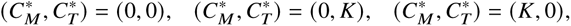

and a line containing infinite number of equilibrium points:

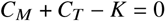

Linear stability analysis shows that (*C_M_*^∗^, *C_T_*^∗^) = (0, 0) has two positive eigenvalues:

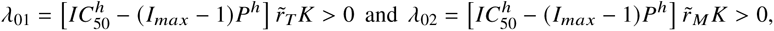

with corresponding eigenvectors

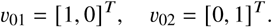

The equilibrium (*C_M_*^∗^, *C_T_*^∗^) = (0, *K*) has eigenvalues:

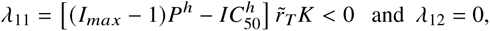

with corresponding eigenvectors

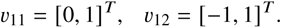

Lastly, the equilibrium (*C_M_*^∗^, *C_T_*^∗^) = (*K*, 0) has two eigenvalues

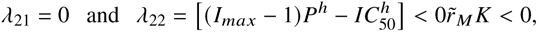

with corresponding eignenvectors

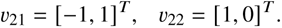

The line of infinite equilibria *C*_*M*_ + *C*_*T*_ − *K* = 0 has two eigenvalues

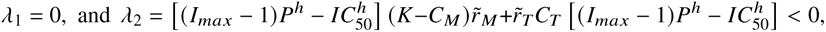

with corresponding eigenvectors

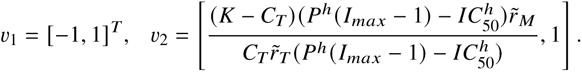

Flow along non-zero eigenvalues is much faster than flow along zero eigenvalues. Therefore, in the fast timescale, cell trajectories are repelled from equilibrium (*C_M_*^∗^, *C_T_*^∗^) = (0, 0) and quickly converge to the neighbourhood of the eigenvector associated with zero eigenvalue. Moreover, the eigenvector associated with zero eigenvalue *υ*_12_ = *υ*_21_ = *υ*_1_ = [−1, 1]^*T*^ collides with the equilibrium line *C*_*M*_ + *C*_*T*_ − *K* = 0, which is the slow manifold. Therefore, in the slow timescales, the flow on the slow manifold has no movement. It implies that the cancer population will eventually converge to the carrying capacity K, but have a different proportion depending on their initial fraction. A simulated vector field under the assumption *K*_*M*_ = *K*_*T*_ = *K* is shown in Figure 23.

**Fig. 23.**
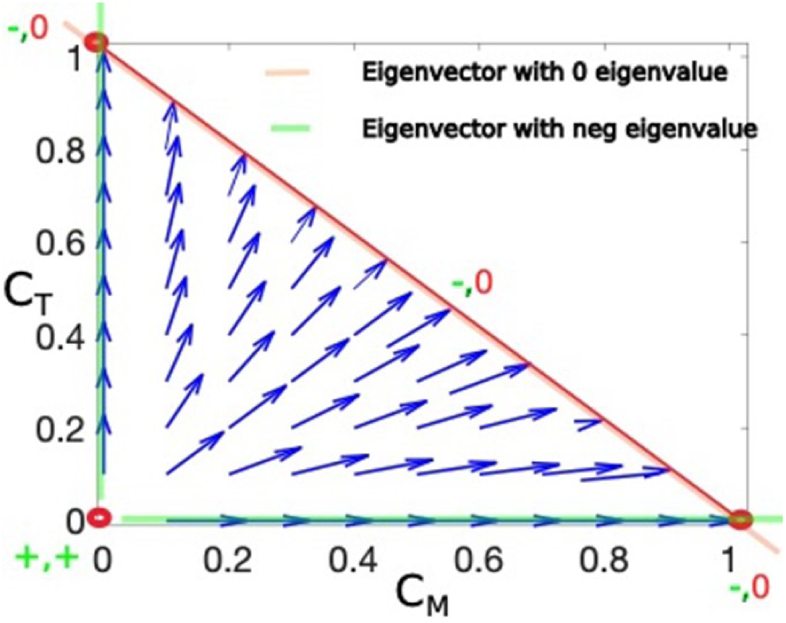
Simulated vector field. Three equilibriums (0, 0), (0, *K*), and (*K*, 0) are denoted in red circles. The line with infinite equilibriums, *C*_*M*_ +*C*_*T*_ −*K* = 0, is plotted in red line. Their eigenvector corresponding to the negative eigenvalues (i.e., *λ*_11_, *λ*_22_, and *λ*_2_) are denoted by the green line. The eigenvector corresponding to the zero eigenvalue is represented in pink line and collapse with the equilibrium line, *C*_*M*_ + *C*_*T*_ − *K* = 0. As a result, the vector field in the blue arrow line points toward the red equilibrium line. Moreover, there is no movement in the red line.

